# Global and local excitation and inhibition shape the dynamics of the cortico-striatal-thalamo-cortical pathway

**DOI:** 10.1101/173658

**Authors:** Anca Rădulescu, Joanna Herron, Caitlin Kennedy, Annalisa Scimemi

**Affiliations:** Department of Mathematics, State University of New York at New Paltz; New York, USA; Department of Biology, State University of Albany at Albany; Albany, USA; Email: or

## Abstract

The cortico-striatal-thalamo-cortical (CSTC) pathway is a brain circuit that controls movement execution, habit formation and reward. Hyperactivity in the CSTC pathway is involved in obsessive compulsive disorder, a neuropsychiatric disorder characterized by the execution of repetitive involuntary movements. The striatum shapes the activity of the CSTC pathway through the coordinated activation of two classes of medium spiny neurons (MSNs) expressing D1 or D2 dopamine receptors. The exact mechanisms by which balanced excitation/inhibition of these cells controls the network dynamics of the CSTC pathway remain unclear. Here we use non-linear modeling of neuronal activity and bifurcation theory to investigate how global and local changes in excitation/inhibition of MSNs regulate the activity of the CSTC pathway. Our findings indicate that a global and proportionate increase in excitation/inhibition pushes the system to states of generalized hyper-activity throughout the entire CSTC pathway. Certain disproportionate changes in global excitation/inhibition trigger network oscillations. Local changes in the excitation/inhibition of MSNs generate specific oscillatory behaviors in MSNs and in the CSTC pathway. These findings indicate that subtle changes in the relative strength of excitation/inhibition of MSNs can powerfully control the network dynamics of the CSTC pathway in ways that are not easily predicted by its synaptic connections.

## Introduction

The cortico-striatal-thalamo-cortical (CSTC) pathway is a neuronal circuit that controls movement selection and initiation, reinforcement and reward [1]. Our current understanding of the wiring diagram of the CSTC pathway, based on functional neuroimaging studies, indicate that this circuit has an equivalent organization in rodents and humans of either sex and age (Fig 1A,B) [9]. In normal conditions, oscillations and synchronous activity in distinct domains of the CSTC pathway are crucial to ensure the execution of habitual actions [2, 3, 41]. In contrast, hyperactivity in specific domains or throughout the circuit are thought to underlie the manifestation of obsessive compulsive disorder (OCD) and Tourette’s syndrome [8, 9]. The basal ganglia nucleus of the striatum plays an important role in regulating the activity of the CSTC pathway through the coordinated interplay of two populations of long-projections GABAergic neurons: the D1 and D2 dopamine receptor expressing medium spiny neurons (D1- and D2-MSNs, respectively) [2, 5]. D1-MSNs project directly to the internal capsule of the globus pallidus (GPi) and the substantia nigra pars reticulata (SNr; Fig 1C), whereas D2-MSNs project indirectly to the SNr by way of intermediate synaptic connections in the external capsule of the globus pallidus (GPe) and the sub-thalamic nucleus (STN; Fig 1C). Activation of D1-MSNs facilitates movement initiation and activation of D2-MSNs inhibits initiation of competing actions [6, 7]. Recent findings indicate that repetitive optogenetic activation of cortico-striatal glutamatergic afferents to the striatum triggers OCD-like behaviors in mice [10], whereas optogenetic stimulation of feed-forward inhibition onto D1- and D2-MSNs relieves OCD-like behaviors in a genetic mouse model of OCD [11]. These findings are important because they suggest that a fine regulation of excitation and inhibition (E/I) onto D1- and D2-MSNs might be crucial to control the onset of hyperactivity in the CSTC pathway [8]. Accordingly, some works suggest that glutamatergic synaptic dysfunction contributes to the etiology of OCD [54], whereas others indicate that this is due to changes in GABAergic inhibition [11, 55, 56, 57]. One way to resolve this conundrum is to determine exactly how changing the weight of excitation and inhibition globally throughout the circuit on local in specific nodes and cell types shapes the network dynamics of the CSTC pathway. Here we address this question using a theoretical approach, based on a mathematical model of the excitatory (glutamatergic) and inhibitory (GABAergic) connections in the CSTC pathway. By using coupled Wilson-Cowan models and bifurcation theory [12], we determine how global and local changes in excitation and inhibition contribute to orchestrate the activity of D1- and D2-MSNs and of the entire CSTC pathway.

**Fig 1.**
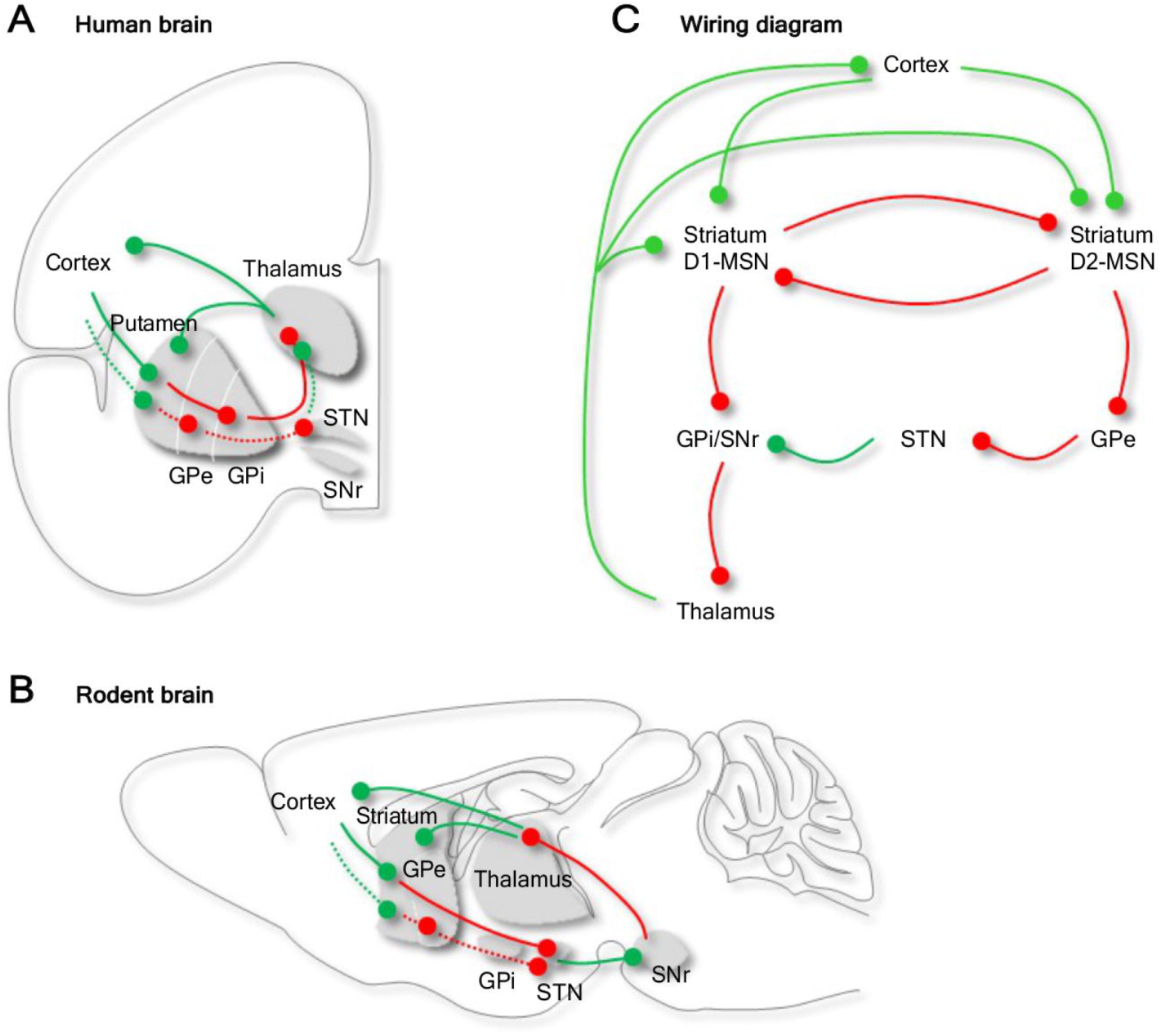
Synaptic connections of the CSTC pathway in humans and rodents. *(A) Anatomical distribution of the synaptic connections of the CSTC pathway in the human brain. The CSTC pathway is a multi*-*synaptic neuronal circuit that connects the cortex*, *striatum and thalamus. The striatum receives glutamatergic inputs (green) from the cortex and the thalamus and sends out GABAergic (red) inputs to the sub-thalamic nucleus. (B) Anatomical distribution of the synaptic connections of the CSTC pathway in the rodent brain. (C) Schematic representations of the synaptic connections in the CSTC pathway. The two most abundant cell types in the striatum*, *D1*- *and D2*-*MSN*, *receive glutamatergic inputs from the cortex and the thalamus and inhibit each other via GABAergic synaptic connections. D1*- *and D2*-*MSNs send GABAergic signals to other basal ganglia nuclei via direct and indirect connections to the substantia nigra (SNr)*, *respectively. Abbreviations: GPe (globus pallidus external part)*, *GPi (globus pallidus internal part)*, *STN (subthalamic nucleus)*, *SNs (substantia nigra pars compacta)*.

## Materials and Methods

We use a system of seven nonlinear equations to describe the major excitatory (glutamatergic) and inhibitory (GABAergic) synaptic connections of the CSTC pathway, thought to be equivalent in rodents and humans of either sex or age (Fig 1) [9]. The general structure of this circuit is conserved across rodents and humans of either sex and age (Fig 1) [9], allowing our findings to be applicable to a variety of contexts and animal species. In this system, each equation provides a quantitative description of the strength of excitation (*c_e_*) and inhibition (*c_i_*) onto each of the seven nodes of the CSTC pathway (Eq 3-10). We then examine the network dynamics of the CSTC pathway using a modeling approach based on the use of analytic equations originally developed by Wilson and Cowan [12] and extensively used in a number of modeling contexts [15, 16, 17]. According to this model, the level of activity in each node (representing the mean activity of a population of neurons) can be described as:

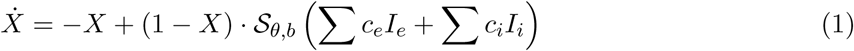

where *X* represents the mean level of activity in the cortex (C), striatum (S), thalamus (T), external and internal capsules of the globus pallidus (E and I, respectively), D1 and D2-MSNs (D1 and D2, respectively). We define *Z* = ∑ *c_e_I_e_* + ∑ *c_i_I_i_* as the weighted sum of all network inputs received by a node *X*. Here *I_e_* and *I_i_* represent the activity in all nodes from which *X* receivs excitatory/inhibitory inputs, whereas *c_e_* and *c_i_* represent average synaptic strengths. The original Wilson-Cowan models obtains theoretical conditions for the synaptic parameters that produce transitions in behavior and then interprets them in a physiological context. Then specific ranges are used to illustrate these conditions and the corresponding bifurcations. Since, for the coupled dynamics, one would expect the relevant dynamic behavior to be encountered in similar parameter ranges as in the original model, we use an extended range that includes all values used in the Wilson-Cowan paper and in subsequent work [12, 14] to describe the synaptic strength values *c_e_* and *c_i_* (0 ≤ *c_e_*, *c_i_* ≤ 40). In this context, values of *c_e_* = 20, *c_i_* = 20 are considered to be indicative of a normal, physiological state. The function 𝓢_*θ*,*b*_ expresses the likelihood that a node responds to the integrated input at time *t* and has sigmoidal shape, which represents the typical physiological neural response to stimuli of increasing strength. Accordingly, a gradual increase in input strength elicits first a low firing response (initial portion of the sigmoidal curve), followed by a window of stimulus strengths that induce profound changes in the firing response (steep portion of the sigmoidal curve), and by a region where the firing rate does not change any more (asymptotic portion of the sigmoidal curve). Eq 1 also incorporates the time constant (identical for all nodes) and the activity history of the node/population. As in the Wilson-Cowan model, in each node, the state *X* = 0 corresponds to a state of low-level background neuronal activity, with small negative values of *X* representing depression of resting activity. Hence the state in which *X* = 0 for all nodes must be a stable steady-state solution for our equations in the absence of external inputs, in order to have physiological significance. In the Wilson-Cowan model, this requirement is fulfilled by setting 𝓢_*θ*,*b*_(0) = 0, therefore considering it of the form:

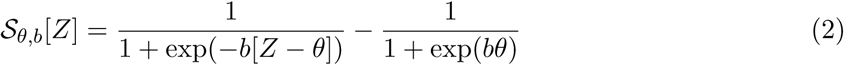

The sigmoidal parameters *θ* and *b* give the value and position of the maximum slope of the sigmoid function in each node. Varying these parameters controls the position and impact of the steep
portion of the curve, but the results of the model remain qualitatively similar regardless of their
specific value (in terms of number and stability of steady states, hysteresis effects, presence of limit cycles, etc.) as long as the general shape of the sigmoidal curve remains the same. Therefore, in this study, we fix the sigmoidal parameters describing excitation (*θ_e_* = 4, *b_e_* = 1.2) and inhibition (*θ_i_* = 2, *b_i_* = 1) to values that are within the range of those originally considered by Wilson-Cowan (1 ≤ *θ_e_*, *θ_i_* ≤ 4; 1 ≤ *b_e_*, *b_i_* ≤ 2 [12]), and consider them as indicative of the physiological control state. Therefore, the state describing the physiological ongoing activity in different nodes of the CSTC pathway is defined as the state where: *c_e_* = 20, *θ_e_* = 4, *b_e_* = 1.2, *c_i_* = 20, *θ_i_* = 2, *b_i_* = 1.

The model exhibits dynamic phenomena typical of a coupled system of non-linear oscillators, which are described in more detail in the Results section. The time constants of all nodes are identical (i.e. all nodes are synchronized, with no aperiodic behavior/chaos). The strength of this modeling framework is that it approaches simultaneously the temporal behavior of seven coupled nodes of the CSTC pathway. However, because the model represents a seven dimensional phase space combined with a multi-dimensional parameter space, it is difficult to perform a global analysis of the system’s dynamics in both phase and parameter spaces. To overcome this potential limitation, one can either perform computer-assisted searches for all types of behaviors that the system can achieve [19], or focus on specific behaviors of interest, which is the approach that we use here. By constructing a sequence of progressively refined models, we can tease apart the effects of gradually finer differences in E/I onto D1- and D2-MSNs in regulating the network dynamics of the CSTC pathway without having to vary more than two key parameters at the same time (i.e. we work in a 2D parameter space). We then shift our attention to the phase space, to study the effects of specific perturbations on the activity of selected nodes.

The mathematical (numerical) analysis of the network dynamics of the system (i.e. the CSTC pathway) is performed as follows. *First*, for fixed parameter values, we analyze the phase space to identify the geometry and position of asymptotic attractors, such as stable equilibria and cycles. The basin of attraction of the stable equilibria is determined by the range of values of the seven-dimensional initial conditions that evolve asymptotically towards that particular attractor. The position of the asymptotic attractors describes the activity of the nodes when they have reached their long-term stable states. Some initial conditions may converge to stable equilibria with higher activity for D1-MSNs than for D2-NSNs. Other initial conditions may converge to stable equilibria with higher activity for D2-MSNs than for D1-MSNs. Some initial conditions may converge to states of perpetual oscillations with characteristic amplitude and duty cycle. *Second*, we analyze how the phase space, with the position and geometry of its attractors, evolves in response to changes in E/I, which we refer to as perturbations. *Third*, we perform a numerical search for bifurcations, which are critical states where the system exhibits sharp transitions between qualitatively different phase space dynamics (e.g. from a non-oscillatory to oscillatory activity). We find co-dimension one bifurcations by studying the sensitivity of the system to changes of one parameter at a time. We find co-dimension two bifurcations by simultaneously changing two parameters at a time. In this study, we specifically analyze how the system responds to independent perturbations of either E or I (and search for co-dimension one bifurcations) or to concurrent perturbations of both E and I (and search for co-dimension two bifurcations). The perturbations are performed on excitatory and inhibitory connections of the entire CSTC pathway or only on those targeting D1- or D2-MSNs.

We use the Matcont package [20, 21] to search specifically for four types of co-dimension one bifurcations: branch points (BPs, or transcritical bifurcations), limit points (LPs, or saddle node bifurcations), Hopf bifurcations (H) and limit point cycles (LPCs, or fold bifurcations). We use this nomenclature in the remainder of the paper for consistency with the Matcont package terminology. BPs occur when two equilibrium curves intersect and simply swap stability. LPs occur when two equilibria collide and vanish. When one of these equilibria is attracting, a drastic rearrangement of the phase space is necessary so that a different attractor may take over the basin of the lost stable equilibrium (hysteresis). In our system, LPs may occur when a perturbation causes a change from high activity state for D1-MSNs to a high activity state for D2-MSNs, even when the system is initiated at the same original state for all variables. H bifurcations occur when an equilibrium suddenly changes stability, giving rise to a limit cycle. We are particularly interested in supercritical Hopf bifurcations, where a stable equilibrium gives birth to a stable cycle that acts as a new asymptotic attractor. At a supercritical H bifurcation, the system may be redirected from a long-term stable state (e.g. high activity for D1-MSNs) to a regime of perpetual oscillations (e.g. between high and low D1-MSN activity). LPCs occur when two cycles collide and disappear (similarly to when two equilibria collide at an LP). If either cycle is an attractor for a set of initial conditions, its loss requires redirecting these solutions towards a different attractor (e.g. a stable equilibrium). An LPC may be, for that reason, a plausible mechanism for cessation of oscillations by small parameter perturbations. In addition to asymptotic attractors, we also search for saddle equilibria and cycles, which only attract solutions originating in lower-dimensional basins (“shallow” basins). For example, at a supercritical Hopf bifurcation, a saddle equilibrium may lose two attracting dimensions to give birth to a saddle cycle with an attracting basin contained in a two-dimensional sub-space of the original seven-variable space. The presence of shallow basins may strongly influence the geometry of other attractors and their own basins, affecting a wide array of trajectories. Shallow basins are hard to identify by randomly choosing initial conditions that eventually converge to a specific saddle point/cycle. In our numerical investigations, we identify saddle equilibria and cycles and their attracting basins whenever possible and we specify when computations become intractable. We consider the system to be in a state of multistability if multiple local attractors co-exist in the phase space, for fixed parameter values, situation which leads to initial conditions that evolve towards different long-term dynamics, depending on the basin of which attractor they belong to. Initial conditions close to the boundary between two basins can easily be pushed towards either attractor by very small perturbations in the initial state of the system (without even changing the system parameters). The ability of the system to swiftly converge to a different steady state from slightly different original states supports the idea that the long-term behavior of the network relies on the E/I balance acting *in conjunction* with the current state of the system. As we identify all of these different behaviors, we point out their relevance for our study. Our models are not independent, but rather developed incrementally, with increasingly subtler phenomena brought under focus with each section.

## Results

### Effect of network-wide changes in E/I

To provide a mathematical representation of the network activity of the CSTC pathway (Fig 1), we use the following system of seven nonlinear equations (see Methods):

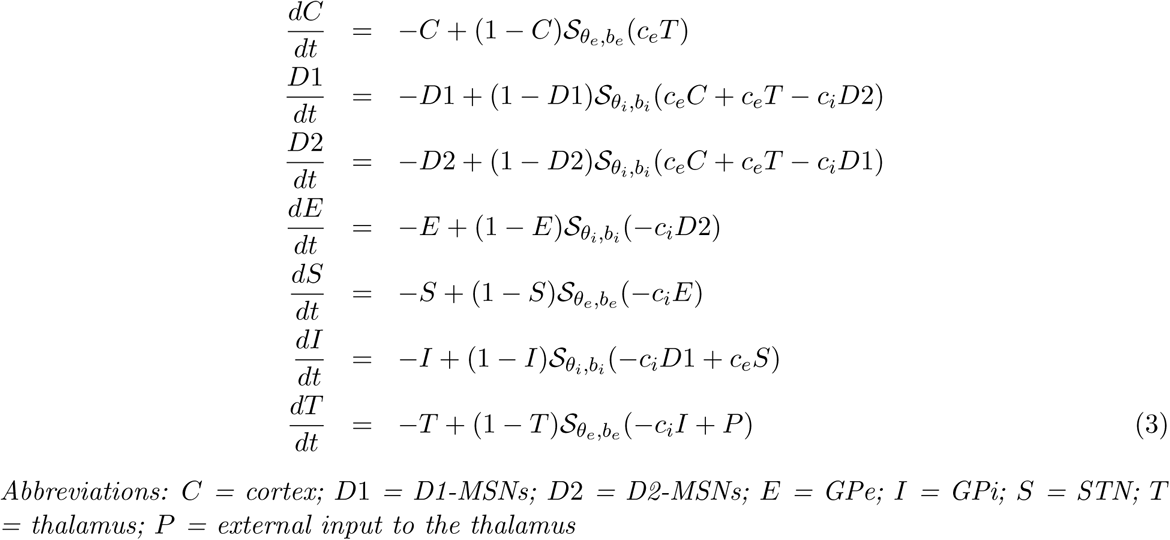

We first assume that all inputs mediating either excitation or inhibition behave identically throughout the entire CSTC pathway and therefore have the same sigmoidal parameters (excitation: *θ_e_* = 4 and *b_e_* = 1.2; inhibition: *θ_i_* = 2 and *b_i_* = 1) and strength (excitation: *c_e_*; inhibition: *c_i_*). We ask how changing the strength of E/I throughout the entire system affects its network dynamics. To address this question, we first set *c_e_* = 8 and *c_i_* = 10 (Fig 2, bottom left). We then vary *c_e_* and *c_i_* progressively until *c_e_* = 30 and *c_i_* = 40 (Fig 2, top right). When *c_e_* and *c_i_* are small, the system initiated at zero stabilizes to an asymptotic attractor of low cortical, striatal and thalamic activity. In contrast, when *c_e_* and *c_i_* are large, the system converges to a state of high activity for the cortex, striatum and thalamus (Fig 2, top right). The transition between these two distinct states with small or large (*c_e_*, *c_i_*) values can occur through different paths, along which *c_e_* is either larger (Fig 2, bottom right) or smaller than *c_i_* (Fig 2, top left). These paths can cross regions of the phase space where the system generates oscillations, with amplitude and duty cycles that depend on the specific values of *c_e_* and *c_i_* (Fig 3). That is to say that the equilibrium curves describing the activity of the cortex, D1- and D2-MSNs and the thalamus show different profiles depending on whether *c_e_* is varied while keeping *c_i_* constant (Fig 3A-D) or whether *c_i_* is varied while keeping *c_e_* constant (Fig 3E-H). When the system reaches a supercritical Hopf bifurcation point, it gains access to a regime of periodic oscillations. In our simulations, network oscillations that arise at the Hopf bifurcation point disappear via a fold bifurcation. Varying only *c_e_* for a specified fixed value of *c_i_*, like varying only *c_i_* for a specified value of *c_e_*, does not significantly affect the position of the Hopf bifurcation point (Fig 3). In contrast, increasing both *c_e_* and *c_i_* alters more efficiently the position of the Hopf bifurcation point and therefore changes the dynamics of the system more profoundly (Fig 3I). Values of (*c_e_*, *c_i_*) that bring the system inside the Hopf loop (blue area in the (*c_e_*, *c_i_*) parameter plane illustrated in Fig 3I) allow the system to show oscillations. Values of (*c_e_*, *c_i_*) that bring the system out of the Hopf loop, prevent it from showing stable oscillations (white and green areas in Fig 3I). The results of this set of simulations show that global changes in either excitation or inhibition control the steady state level of activity of the CSTC pathway. In contrast, concurrent changes in global E/I control the stability of the system, allowing it to show network oscillations with specific amplitudes and duty cycles.

**Fig 2.**
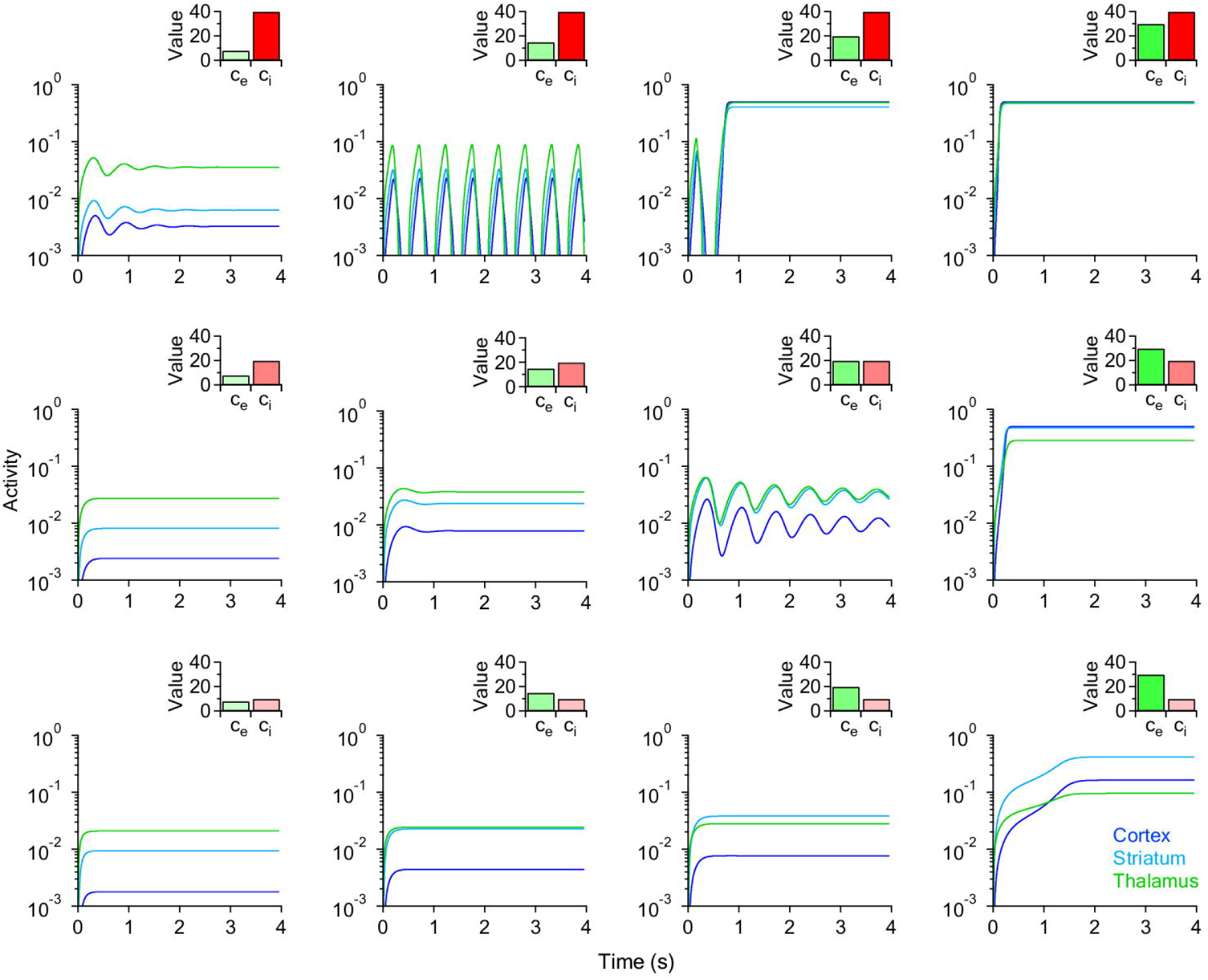
Evolution of the temporal behavior of the CSTC pathway in response to changes in global excitation (*c_e_*) and inhibition (*c_i_*). *Each curve represents the temporal evolution in the activity of the cortex (blue)*, *striatum (cyan) and thalamus (green)*. *The insets show the values of c_e_ and c_i_ used in each set of simulations (c_e_* = 8, 15, 20, 30 *and c_i_* = 10, 20, 40*). Excitation increases from left to right in each row. Inhibition increases from bottom to top in each column. Other parameters are fixed as follows: τ_e_* = 4, *b_e_* = 1.2, *τ_i_* = 2, *b_i_* = 1, *P* = 1.

**Fig 3.**
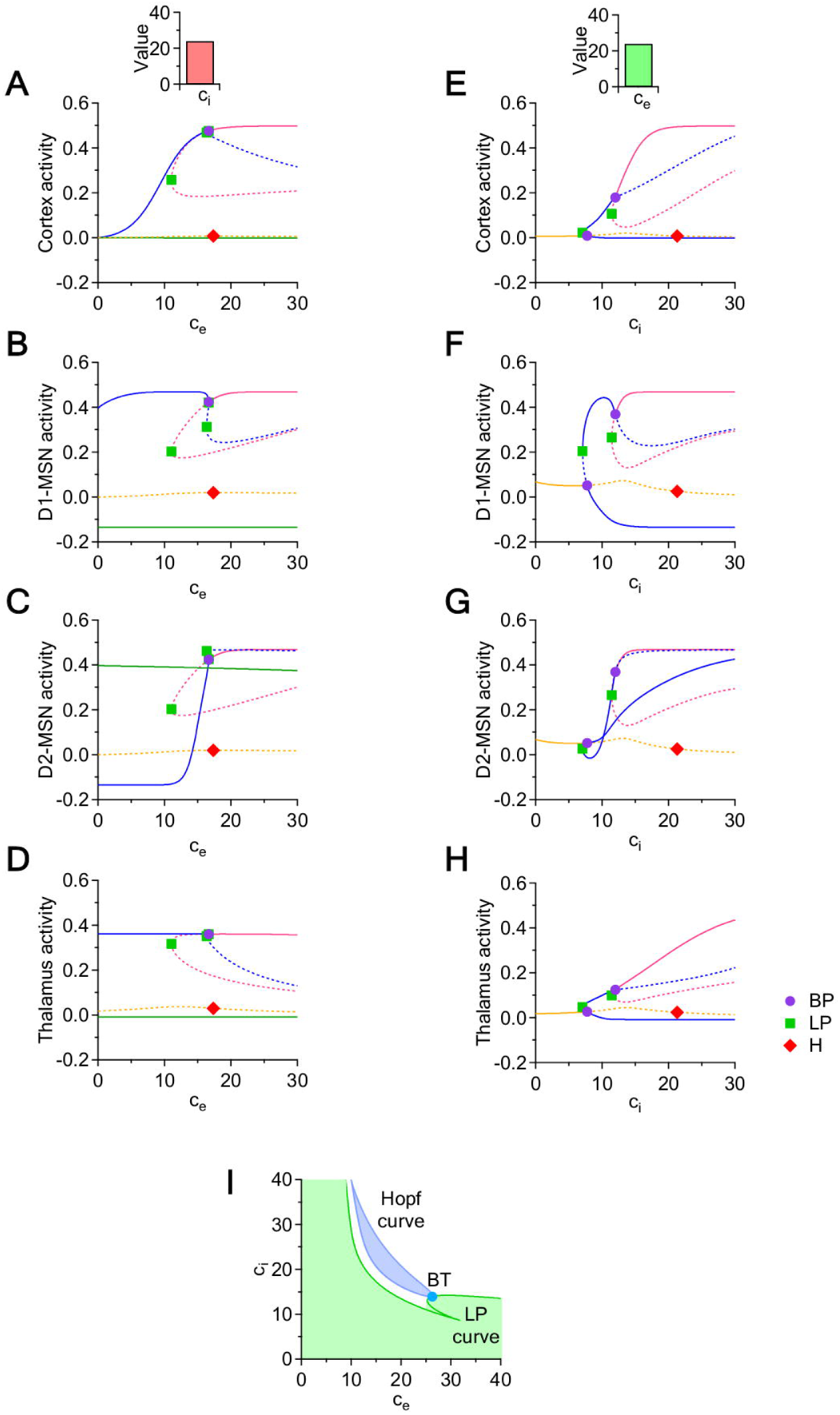
Bifurcation diagrams with respect to *c_e_* and *c_i_* and bifurcation curves in the (*c_e_*, *c_i_*) parameter plane. *(A*-*H) Each panel shows multiple equilibrium curves for the cortex (A*,*E)*, *D1*-*MSNs (B*,*F), D2*-*MSNs (C*,*G) and the thalamus (D*,*H). Panels (A*-*D) are obtained by varying only c_e_ while keeping c_i_* = 24. *Panels (E*-*F) are obtained by varying c_i_ while keeping c_e_* = 24. *The equilibrium curves show that the system has multistability windows for most values of c_e_ and c_i_. In each panel*, *the symbols represent codimension one bifurcation points: branch point (BP*, *purple circle)*, *limit point (LP*, *green square)*, *Hopf point (H*, *red diamond). (I) Local bifurcation curves in the (c_e_*, *c_i_) parameter plane. The blue loop represents a Hopf bifurcation curve*, *which delimits a regime of oscillations (blue area) from a regime of convergence to a steady state (white and green areas). The green curve is an LP curve*, *along which two equilibria (in the region to the right of the green curve) collide and disappear (in the region to the left of the green curve). The Hopf and the LP curves touch at a Bogdanov-Takens (BT) codimension two bifurcation point.*

### Effect of local changes in E/I onto D1- and D2 -MSNs

We next ask how local changes in E/I specifically targeted onto D1- and D2-MSNs affect the network dynamics of the CSTC pathway. To address this question, we add two new parameters to the system of seven nonlinear equations, which we name
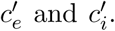
We begin our analysis by altering the strength of excitation onto D1- and D2-MSNs. To do so, we change
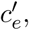while keeping
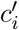
fixed (Section and Fig 4A-D). Then, we alter the strength of inhibition onto D1- and D2-MSNs. This is accomplished by changing
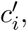
while keeping
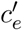
fixed (Section and Fig 4E-H). As in the previous model, we analyze the evolution of the system’s asymptotic behavior in the cortex, D1- and D2-MSNs and the thalamus.

**Fig 4.**
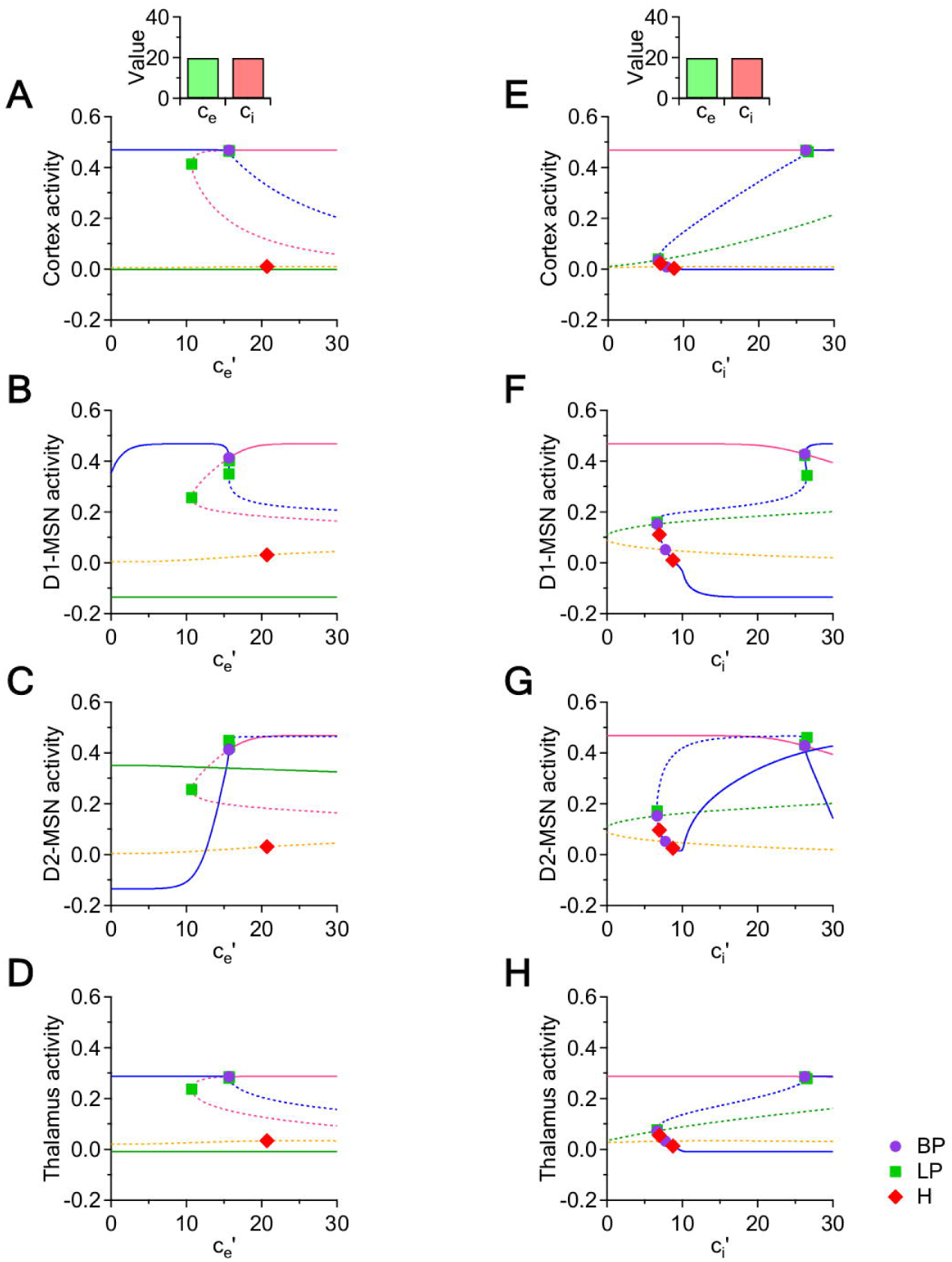
Bifurcation diagrams with respect to
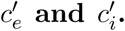. *(A*-*D) In each panel*, *we test how changing the strength of local excitation onto D1*- *and D2-MSNs (i.e.*
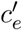
*) affects the activity of the cortex*, *of the two populations of MSNs and the thalamus. The strength of excitation and inhibition in the rest of the system is set to c_e_* = *c_i_* = 20 *(top inset). (E*-*H) In each panel*, *we analyze how changing the strength of inhibition onto D1*- *and D2*-*MSNs (i.e.*
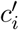
*) affects the activity of the cortex*, *MSNs and the thalamus. The strength of excitation and inhibition in the rest of the system is set to c_e_* = *c_i_* = 20.

### Effect of local changes in excitation onto D1- and D2-MSNs

We use the following system of equations to determine the effect of changing the strength of local excitation onto D1- and D2-MSNs (i.e.
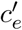).

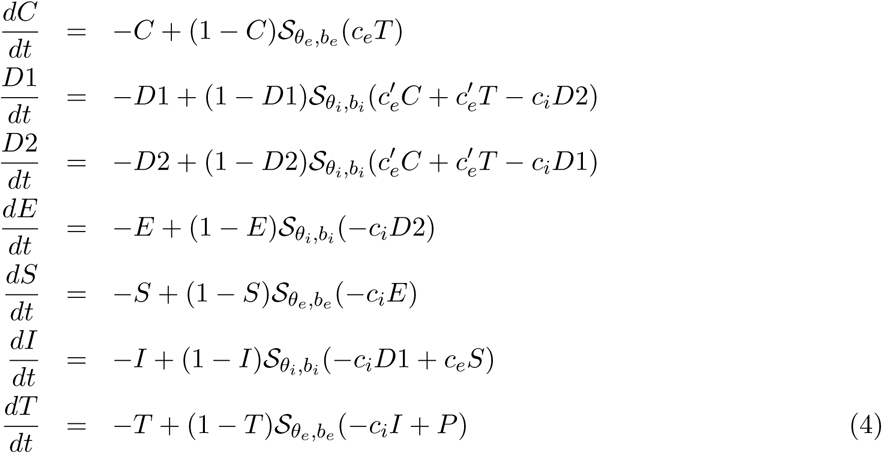

The most notable feature in this set of simulations is that the system shows a bi-stable behavior along the entire range of tested values for
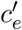
(Fig 4). When
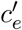
is small (5 <
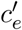
< 10), the system has two stable equilibrium curves (blue and green curves) and one saddle equilibrium curve (orange curve). If all variables are initiated at rest (zero initial conditions), the system converges to the orange equilibrium; if the rest initial conditions are slightly perturbed in the D1-MSN component, then the system converges to the blue equilibrium (Fig 4B); if they are perturbed in the D2-MSN component, then the system converges to the the green equilibrium (Fig 4C). The position of all these equilibrium curves remains approximately constant, allowing the system to remain either in a stable state of high activity for the cortex, D1-MSNs and thalamus and low activity for D2-MSNs, or in a stable state of low activity for the cortex, D1-MSNs and thalamus and high activity for D2-MSNs. As
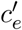
increases (10 <
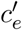
< 15), the stable (green) and saddle (orange) branches do not change appreciably, while the stable (blue) branch increases steadily in its D2-MSNs component. At the *BP*, when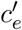
~ 15, the upper part of the pink branch takes over the stability from the blue branch, and maintains, for values of
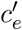
higher than *BP*, the asymptotic state of high activity in all four regions of the system, attracting all initial conditions with higher activity of D1- versus D2-MSNs. A decrease in the initial activity of D1- versus D2-MSNs can lead the system to converge towards the other stable state (green), of low activity for the cortex, D1-MSNs and thalamus and high activity for D2-MSNs (green). Hence different types of unbalanced activity of D1- versus D2-MSNs can bring the system into one or the other attraction basins. This new bistable asymptotic regime survives without state transitions for all values of
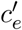
> 15. A perfect balance in the activity of D1- and D2-MSNs allows the system to reach a saddle equilibrium curve (orange curve) for all values of
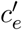
. This saddle equilibrium curve corresponds to a low activity state in the cortex, striatum and thalamus, which persists until the Hopf bifurcation point, where the system gains a saddle cycle.

### Effect of local changes in inhibition onto D1- and D2-MSNs

We use the following system of equations to determine the effect of changing the strength of local inhibition onto D1- and D2-MSNs (i.e.
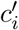).

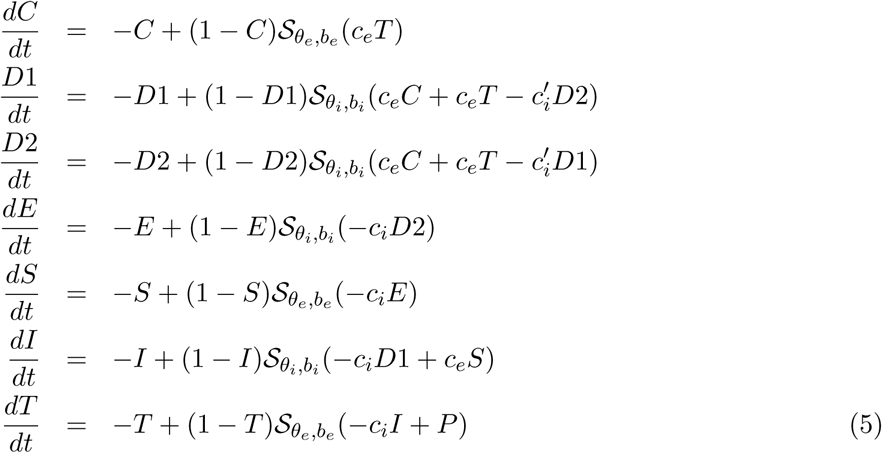

A first notable feature in these simulations is that multiple equilibrium curves remain similar along the entire range of values for
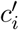
(Fig 4E-H). When 0 <
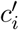
< 7, the system converges to a steady state characterized by high activity in the cortex, D1- and D2-MSNs and thalamus (pink curve). A second notable feature is the presence of stable limit cycles. When
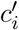
~ 7, the system undergoes a fold bifurcation and generates a stable cycle (not shown in Fig 4), which later collides with the blue equilbrium curve. More precisely, this cycle coexists with the blue equilibrium branch within a narrow window of
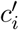
values (7 <
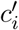
< 9). Then it disappears through a Hopf bifurcation when
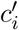
~ 9, changing stability of the blue equilibrium curve from a saddle to an attracting equilibrium. When
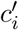
increases beyond the Hopf bifurcation value (
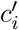
> 9), the system reaches a regime with two stable equilibria (blue and pink curves). One stable branch that emerges from the Hopf point (blue curve) corresponds to a state of high activity in the cortex, thalamus and D2-MSNs, coupled with low activity in D1-MSNs. The second branch (pink curve) corresponds to state of high activity in all brain regions, which preserves high activity levels for both D1- and D2-MSNs until the following bifurcation point (
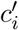
= 26). Here the blue branch loses stability and leads to a dramatic drop in in the activity of D2-MSNs. When
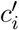
> 26, the D2-MSN component of the blue equilibrium curve decays rapidly, while all other system components retain high activity levels. These findings indicate that changing the strength of local inhibition onto D1- and D2-MSNs allows these cells to show not only concurrent or non-concurrent high activity levels, but also stable oscillations between these states.

### Effect of local changes in the relative E/I of D1- versus D2 -MSNs

We next ask how the activity of the system changes when altering the relative excitation or the relative inhibition of D1- and D2-MSNs. To do this, we keep constant the excitation and inhibition in the rest of the CSTC pathway (*c_e_* = *c_i_*) and then introduce new variables to vary either: (1) the relative strength of excitation onto D1-MSNs (*c*_*e*1_) and D2-MSNs (*c*_*e*2_) or (2) the relative strength of inhibition onto D1-MSNs (*c*_*i*1_) and D2-MSNs (*c*_*i*2_). In both cases, we work with a two-dimensional parameter space, as described in the Methods section.

### Effect of local changes in the relative excitation of D1- versus D2-MSNs

We use the following system of equations to determine the effect of changing the relative strength of excitation onto D1- and D2-MSNs.

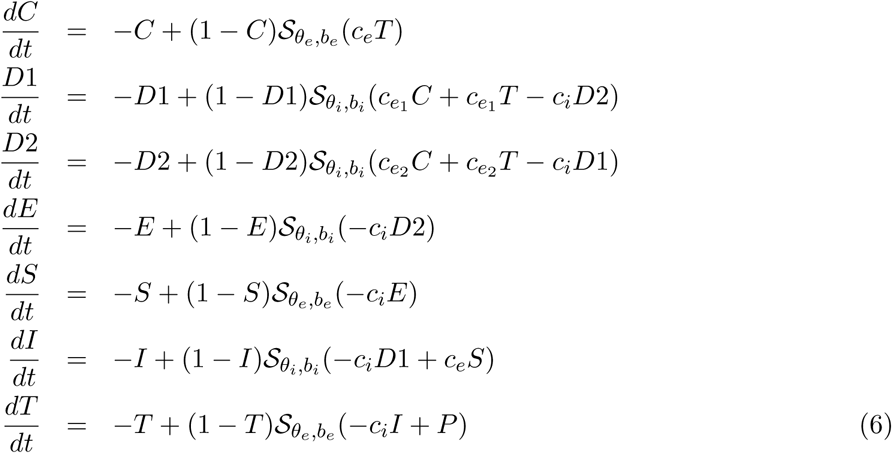

We fix *c_e_* = *c_i_* = 20, and perform two sets of simulations, in which we vary excitation onto D1-MSNs progressively (*c*_*e*_1__), while excitation onto D2-MSNs is set at *c*_*e*_2__ = 10 (Fig 5A-D) and *c*_*e*_2__ = 20 (Fig 5E-H). Inhibition onto D1- and D2-MSNs matches the level of inhibition in the rest of the system and therefore *c*_*i*_1__ = *c*_*i*_2__ = *c_i_*. We let *c*_*e*_1__ vary between *c*_*e*_1__ = 0 and *c*_*e*_1__ = 40. Both sets of simulations identify two stable branches in the equilibrium curves with respect to *c*_*e*_1__ : the pink branch born via the LP at *c*_*e*_1__ ~ 3 and the orange curve that remains constant for the whole *c*_*e*_1__ range (Fig 5A and B). All other branches that we identified numerically are unstable saddles, and the supercritical Hopf bifurcation marked along the pink curve creates saddle cycles with shallow basins (not captured in the subsequent phase planes). Phase plane plots confirm that, when *c*_*e*_2__ = 10, increasing the strength of excitation onto D1-MSNs from *c*_*e*_1__ = 0 to *c*_*e*_1__ = 30 allows the system to transition from mono-stability when 0 < *c*_*e*_1__ < 3 (a unique and stable node of low activity for D1-MSNs and high activity for D2-MSNs; Fig 5C) to bi-stability when *c*_*e*_1__ > 3 (co-existence of this existing node with a new stable node of high activity for D1-MSNs; Fig 5D). As mentioned above, this transition occurs through an LP bifurcation (Fig 5A-B). Similarly, when setting *c*_*e*_2__ = 20, th system has access to a stable steady state with high D2-MSNs activity levels throughout the range of *c*_*e*_1__ (Fig 5E-F). The system exhibits a similar phase transition between mono- and bi-stability, but in this case the transition appears for larger values of *c*_*e*_1__ (*c*_*e*_1__ ~ 15). When *c*_*e*_2__ < 15, the only possible activity level of D1-MSNs is low (Fig 5G). When *c*_*e*_2__ > 15, the system gains access to an alternative stable state where both D1- and D2-MSNs have high activity levels (Fig 5H). Taken together, these findings indicate that increasing excitation of D1-MSNs generates bistability irrespective of the levels of excitation of D2-MSNs. When excitation of D2-MSNs is low, the activity level of D1-MSNs is opposite to the activity level of D2-MSNs: high activity in D1-MSNs causes low activity in D2-MSNs and vice versa (Fig 5C-D). When excitation of D2-MSNs is high, the activity level of D2-MSNs remains high regardless of the activity level of D1-MSNs. Therefore, high levels of excitation of D2-MSNs can be coupled with either high or low activity levels of D1-MSNs (Fig 5G-H).

**Fig 5.**
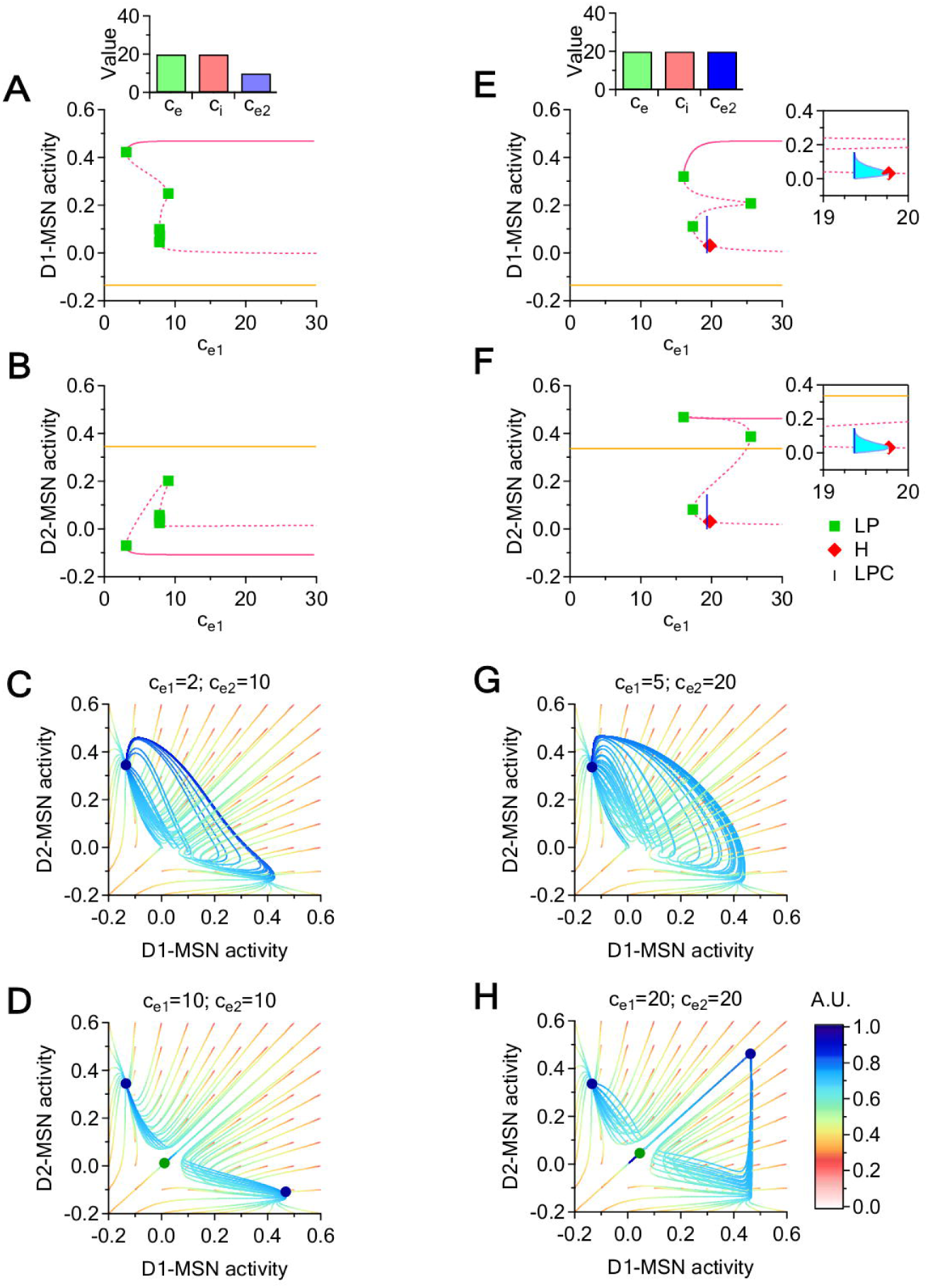
Bifurcations diagrams with respect to *c*_*e*_1__ for increasing values of *c*_*e*_2__, and corresponding phase plane (*D*1, *D*2) slices. *(A*-*B) Bifurcation diagrams showing D1*-*(A) and D2*-*MSNs (B) coordinate projections for increasing values of c*_*e*_1__, *when c*_*e*_2__ = 10. *Each panel shows the position of stable (solid line) and unstable equilibrium curves (dotted lines). The diagrams include LPs (green squares)*, *supercritical H bifurcation points (red diamonds) and LPCs (blue vertical line). (C*-*D) Phase plane dynamics for c*_*e*_2__ = 10. *The panels show phase plane slices*, *in which each curve represents the projection of a seven dimensional trajectory to the (D*1, *D*2*) plane. Each phase plane plot uses a grid of initial conditions of resolution 0.1 within the (D*1, *D*2*) plane. All temporal trajectories evolve from red to cyan. Stable equilibria are shown as blue circles and saddle points are shown as green circles. (E*-*F) As in (A*-*B)*, *for c*_*e*_2__ = 20. *The insets show saddle cycles born at supercritical H bifurcations (cyan). The cycles stop at the blue vertical line representing an LPC bifurcation*, *at which the cycle branch born from the H point collides with another cycle branch (not shown) and disappears. (G*-*H) Phase plane dynamics for c*_*e*_2__ = 20.

### Effect of local changes in the relative inhibition of D1- versus D2-MSNs

We use the following system to study the effect of altering the relative strength of inhibition onto D1- and D2-MSNs:

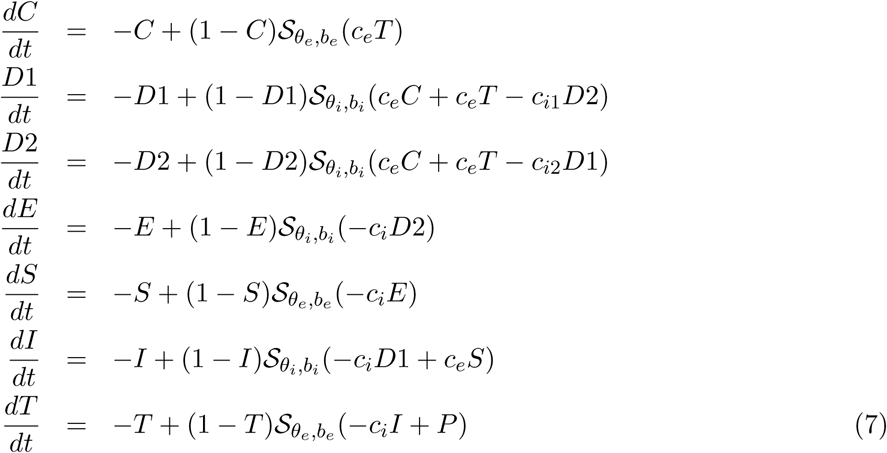

Similarly to our analysis of local excitation, we fix *c_e_* = *c_i_* = 20. We then perform two sets of simulations, in which we vary inhibition onto D1-MSNs (*c*_*i*_1__) while keeping inhibition onto D2-MSNs fixed at *c*_*i*_2__ = 7 (Fig 6A-E) and *c*_*i*_2__ = 20 (Fig 6F-J). Excitation onto D1- and D2-MSNs matches the level of excitation in the rest of the system and therefore *c*_*e*_1__ = *c*_*e*_2__ = *c_e_*. We let *c*_*i*_1__ vary between *c*_*i*_1__ = 0 and *c*_*i*_1__ = 30. The bifurcation diagrams show hysteresis regardless of the specific value of *c*_*i*_2__ (Fig 6A-B, F-G). We first considered the case of *c*_*i*_2__ = 7 (Fig 6A-B). For *c*_*i*_1__ < 10, the system has a unique stable regime, characterized by high activity in both D1- and D2-MSNs, regime which lasts for a wide range of *c*_*i*_1__ values (0 < *c*_*i*_1__ < 26). For *c*_*i*_1__ > 10, a second stable regime appears, characterized by low activity in both D1- and D2-MSNs. We then considered the case of *c*_*i*_2__ = 20 (Fig 6F-G). The high D1- and D2-MSN activity attracting regime is also stable for a wide range of *c*_*i*_1__ values (0 < *c*_*i*_1__ < 26). In fact, this portion of the equilibrium curve is almost identical to its counterpart for the case of *c*_*i*_2__ = 7. However, in our current case, the second stable regime emerges sooner, when *c*_*i*_1__ > 2.2, and leads to low asymptotic activity in D1-MSNs and high activity in D2-MSNs. Significant differences in the behavior of D1- and D2-MSNs are evident at a more detailed investigation moving along the equilibrium curve. When *c*_*i*_2__ = 7, the top branch of the equilibrium curve loses stability at the LP (*c*_*i*_1__ ~ 26.2) until it reaches the supercritical Hopf bifurcation point H (*c*_*i*_1__ ~ 10), where the stable cycle born at the LPC (*c*_*i*_1__ ~ 7.58) collides with the saddle equillibrium and confers it full stability until the next LP (*c*_*i*_1__ ~ 20). After a brief unstable interval (20 < *c*_*i*_1__ < 20.7), a new stable equilibrium is reached. Therefore, under these conditions, three attractors coexist in the narrow parameter window [20, 20.7]. When *c*_*i*_2__ = 20, there is a wide range of *c*_*i*_1__ values that can lead to a regime equivalent to bi-stability (2.2 < *c*_*i*_2__ < 26.2), in the sense that the saddle equilibria and cycle that coexist with these attractors have shallow attraction basins, and are thus unlikely to be reached by generic initial conditions.

**Fig 6.**
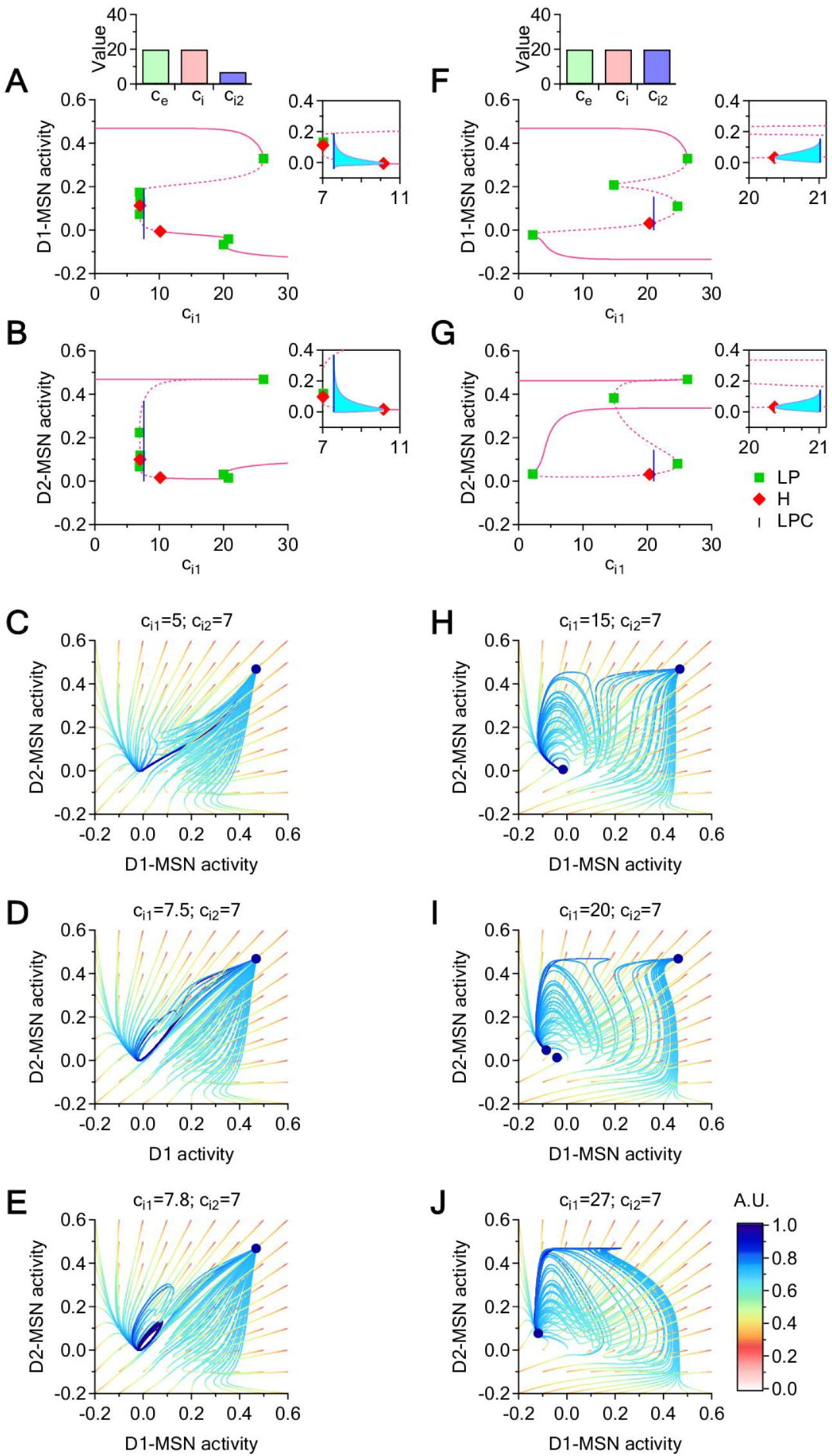
Bifurcation diagrams with respect to *c*_*i*_1__ and phase space trajectories for *c*_*i*_2__ = 7 and different values of *c*_*i*_1__. *(A*-*B) Bifurcation diagrams showing the D*1 *and D*2 *coordinate projections for increasing values of c*_*i*_1__, *when c*_*i*_2__ = 7. *The equilibrium curve (pink) displays six LPs (green squares) and two H points (red diamonds). The insets show the cycles born at the H points (cyan). At the leftmost supercritical H point (c*_*i*_1__ = 7), *the saddle equilibrium changes to a saddle cycle. At the rightmost supercritical H point (c*_*i*_1__ = 10.2), *the stable equilibrium changes to a stable cycle*, *which survives until the LPC at c*_*i*_1__ = 7.6 *(blue line). (C*-*E) Representation of the seven dimensional trajectories via their projections onto the 2*-*dimensional* (*D*1, *D*2) *plane*, *for a grid of initial conditions within the* (*D*1, *D*2) *slice. Trajectories are evolving in time from red to blue. The position of the attractors (equilibria and cycles) is marked with a dark blue circle. (F*-*G) As in (A*-*B)*, *for c*_*i*_2__ = 20. *(H*-*J) As in (C*-*E)*, *for different values of c*_*i*_1__.

The phase diagrams in Fig 6C-E,H-J illustrate some of these behaviors in more detail for *c*_*i*_2__ = 7, in a (*D*1, *D*2) phase space slice. When *c*_*i*_1__ < 7.58, the system has a unique attracting equilibrium with high D1- and D2-MSN activity (Fig 6C-D). When *c*_*i*_1__ = 7.58, it crosses the LPC bifurcation, leading to the birth of a stable cycle and the onset of network oscillations which co-exists with the original attracting equilibrium (Fig 6E). At the Hopf bifurcation point (*H* ~ 10.15), the cycle collapses into a low D1- and D2-MSN activity equilibrium, which continues to coexist with the original high D1- and D2-MSN equilibrium (Fig 6H). Through a short hysteretic window, flanked by *LP* = 19.97 and *LP* = 20.77, a new low equilibrium of low D1- and D2-MSN activity is born (Fig 6I) and replaces the pre-existing low D1- and D2-MSN activity equilibrium (Fig 6J).

In order to illustrate more comprehensively the simultaneous dependence of the system’s behavior on both inhibition levels *c*_*i*_1__ and *c*_*i*_2__, we plot bifurcation curves in the (*c*_*i*_1__, *c*_*i*_2__) parameter plane (Fig 7A). We focus in a region around a Zero-Hopf co-dimension two bifurcation point (ZH), where three bifurcation curves intersect, separating three parameter regions that identify three distinct dynamic behaviors, illustrated in Fig 7B-D. The green region, analyzed in Fig 7B, is characterized by the presence of two stable equilibria, one with high D1- and D2-MSN activity, the other with low D1- and D2-MSN activity. The blue region, analyzed in Fig 7C, is characterized by the formation of a stable cycle when entering from the green region by crossing the H (green) curve. The cycle vanishes via a fold bifurcation when crossing the LPC (blue) curve, so that in the pink region only the stable equilibrium of high D1- and D2-MSN activity survives (Fig 7D). Finally, one can re-enter the green region by crossing the LP curve (pink), with formation of two new equilibria (a node and a saddle), bringing the system back to the initial bistability (Fig 7B).

**Fig 7.**
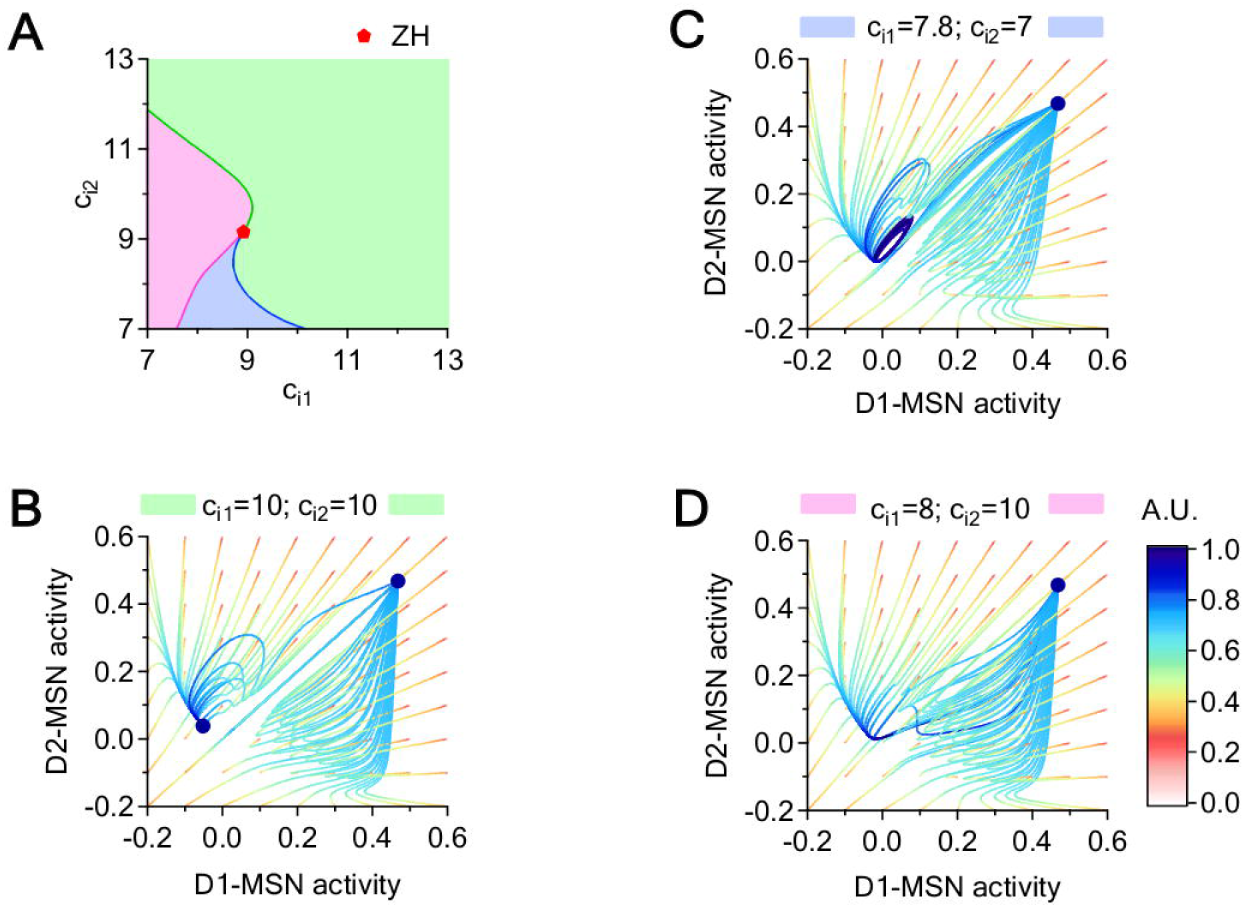
Local bifurcation curves in the (*c*_*i*_1__, *c*_*i*_2__) parameter plane and attracting regimes for different (*c*_*i*_1__, *c*_*i*_2__) pairs. *(A) Intersection of H (blue)*, *LP (green) and LPC (pink) curves at a Zero Hopf (ZH) point. The three different regimes delimited by these curves are color*-*coded in pink*, *green and blue. (B*-*D) Each panel illustrates a generic phase plane slice for one of the three parameter regions*.

Taken together, these findings indicate that increasing inhibition onto D1-MSNs when inhibition onto D2-MSNs is low introduces a bistable behavior where both cell types show either high or low levels of activity. In contrast, increasing inhibition onto D1-MSNs when inhibition onto D2-MSNs is high allows to change the activity level only in D1-MSNs, while the activity of D2-MSNs remains high. In the previous section we saw that altering the relative excitation of D1- and D2-MSNs allows the system to transition through dynamics regimes with limit cycles and saddle cycles, which attract only a shallow subspace of initial conditions. Here we show that altering the relative inhibition of D1- and D2-MSNs pushes the system through dynamics windows with stable cycles with attraction basins of significant size. Therefore, altering the relative inhibition of D1- and D2-MSNs provides a more powerful mechanism than altering the relative excitation of these cells, which can push the system into a regime of stable oscillations.

### Effect of local and coupled changes in the E/I of D1- and D2 -MSNs

In the previous sections, we considered simplified scenarios where we changed either excitation or inhibition onto D1-MSNs, while keeping excitation or inhibition onto D2-MSNs constant at arbitrary values. Here we consider the more complex scenario in which the two effects are coupled. *First*, we change the magnitude of either local excitation or local inhibition proportionately onto D1- and D2-MSNs (using the parameter *h*). *Second*, we scale these effects differently in D1- and D2-MSNs (using the parameters *a* and *b*).

### Effect of local and coupled changes in the excitation of D1- and D2 -MSNs

We use the following system to study the effect of a coupled modulation of excitation onto D1- and D2-MSNs:

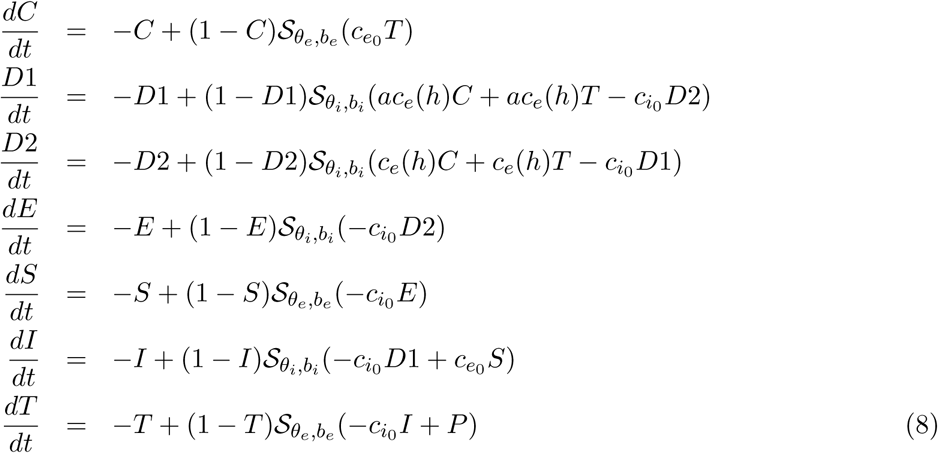

We set the global level of excitation and inhibition in the CSTC pathway to *c*_*e*_0__ = *c*_*i*_0__ = 15, and then we vary the strength of excitatory connections onto D1- and D2-MSNs by changing the *h* parameter *h*. We consider *c_e_*(*h*) = *c*_*e*_0__ + (1 − *e*^−*h*^)*c*_*e*_0__, so that increasing *h* allows us to increase the strength of local excitatory connections *c_e_*(*h*) in both D1 and D2-MSNs in a nonlinear fashion, from a minimum value of *c_e_*(0) = *c*_*e*_0__ to a maximum value of *c_e_*(∞) = 2*c*_*e*_0__. We analyze these effects at two different values of *a*. By increasing *a*, we scale the effects induced by varying *h* differently in D1- and D2-MSNs.

Here we analyze two scenarios in which we vary *h*: when *a* = 1(Fig 8A-D) and when *a* > 1 (Fig 8E-H). We plot a series of bifurcation curves obtaine by varying *h* over an extended (and partly unbiological) range (− 1 < *h* < 2), to visualize the origin of the bifurcation phenomena. Consistent with our previous model (Fig 5), increasing *h* when *a* = 1 moves the system between two bi-stability windows, where a low D1- and high D2-MSN activity state (green curve) coexists with a high D1- and low D2-MSN activity state (orange curve; Fig 8A-D). Increasing *h* moves the system between two distinct bi-stability windows, where the low D1- and high D2-MSN activity state (green curve) coexists with a high D1- and D2-MSN activity state (orange curve). As expected, these observed results reproduce, in part, the behaviors observed when changing excitation onto D1- and D2-MSNs independently (Fig 5). This is to be expected, since this model introduces only a re-parametrization of the excitation strengths from the (*c*_*e*_1__, *c*_*e*_2__) parameter plane to the new (*h*, *a*) parameter plane. Similar findings are observed for a larger scaling factor *a* = 1.2 (Fig 8,E-H). These findings suggest that local changes in excitation onto D1- and D2-MSNs can alter the level of activity in these cells regarless of the fact that they may be more or less pronounced in one of the two cell populations.

**Fig 8.**
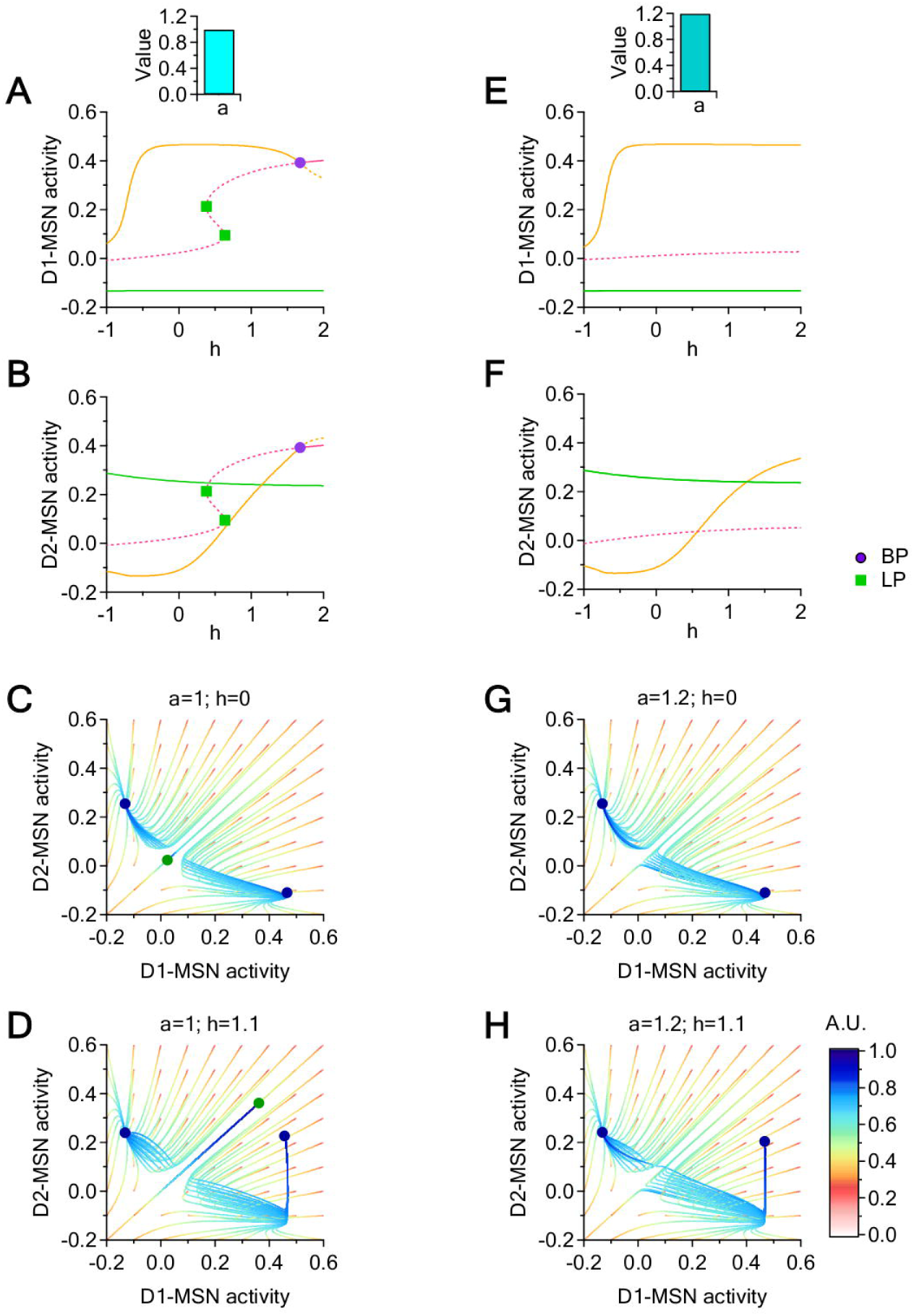
Dynamic transitions for *a* = 1 and *a* = 1.2. *(A*-*B) Bifurcation diagrams showing the D1*- *and D2*-*MSN variable coordinates with respect to h*, *when a* = 1*. (C*-*D) Trajectories in a* (*D*1, *D*2) *phase space slice for h* = 0 *and h* = 1.1, *when a* = 1. *(E*-*F) As in (A*-*B)*, *for a* = 1.2*. (G*-*H) As in (C*-*D)*, *for a* = 1.2*. Fixed parameters: c*_*e*0_ = *c*_*i*_0__ = 15.

### Effect of local and coupled changes in the inhibition of D1- and D2 -MSNs

We use a similar approach to determine the network effects of local changes in inhibition. To do this, we use the following system of equations:

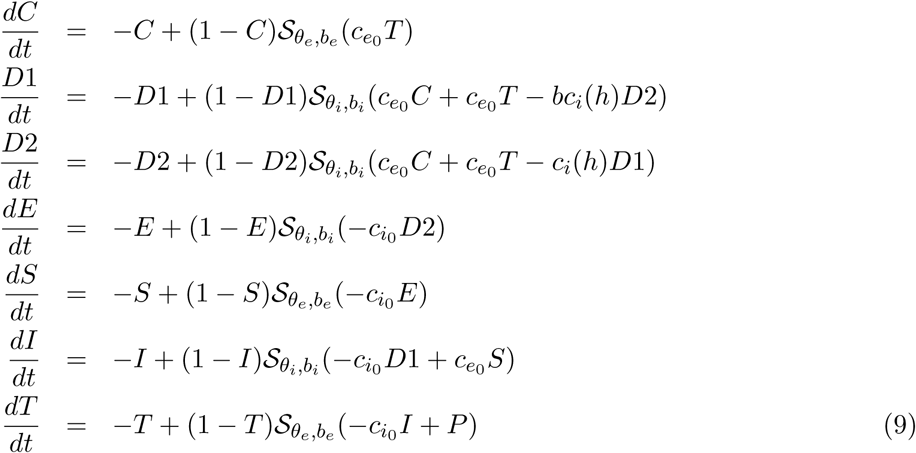

We set the global level of excitation and inhibition in the CSTC pathway to *c*_*e*_0__ = *c*_*i*_0__ = 20, and then we vary the strength of inhibitory connections onto D1- and D2-MSNs by changing the parameter *h*. By changing the parameter *b*, we scale the magnitude of these effects in D1- and D2-MSNs. In this *h* case, we set: *c_i_*(*h*) = *c*_*i*_0__ − (1 − *e*^−*h*^)*c*_*i*_0__, so that by increasing *h*, we decrease the inhibitory synaptic strengths *c_i_*(*h*) in a nonlinear way, from a maximum value of *c_i_*(0) = *c*_*i*_0__ to a minimum value of *c_i_*(1) = 0. We then use the parameter *b* to allow changes in *h* to differentially affect inhibition onto D1- and D2-MSNs. We analyze three scenarios in which we vary *h* when *b* = 0.8, *b* = 1 and *b* = 1.2 (Fig 9A-D, E-J and K-O, respectively). When *b* = 0.8, progressively increasing *h* allows the system to transition from a bi-stable regime of high D1- and D2-MSN activity coexisting with low D1- and high D2-MSN activity, to a regime of high D1- and D2-MSN stable activity, via an LP bifurcation (Fig 9A-D). When *b* = 1, changes in *h* allow the system to transition between the same two regimes described for *b* = 0.8 (i.e. bistability when *h* is small, mono-stability when *h* is large; Fig 9E-J). In addition, when *b* = 1, increasing *h* can trigger oscillations via two supercritical Hopf bifurcations (Fig 9F-G). The first Hopf point (at *h* ~ 0.04) gives birth to a saddle cycle (Fig 9E) which survives until *h* ~ 0.17 and disappears via a LPC. This saddle cycle remains symmetric for the whole duration of its life (i.e. *D*1 = *D*2), and has an attraction basin that also lies in the sub-space *D*1 = *D*2 (Fig 9E). The second Hopf point (at *h* = 0.82) moves the system from bi-stability (Fig 9H) into stable oscillations (Fig 9I), which disappear at *h* ~ 0.92 via a LPC. This cycle attractor coexist throughout its life with a high D1- and D2-MSN stable equilibrium (Fig 9I), before disappearing for larger values of *h* (Fig 9J). Setting *b* = 1.2 generates qualitatively similar results, only in a more efficient way. Therefore, under these conditions, a progressively larger reduction of inhibition onto D1- than D2-MSNs evoked by increasing *h* allows the system to transition from bi-stability to small stable oscillations to a single high-activity equilibrium (Fig 9K-O).

**Fig 9.**
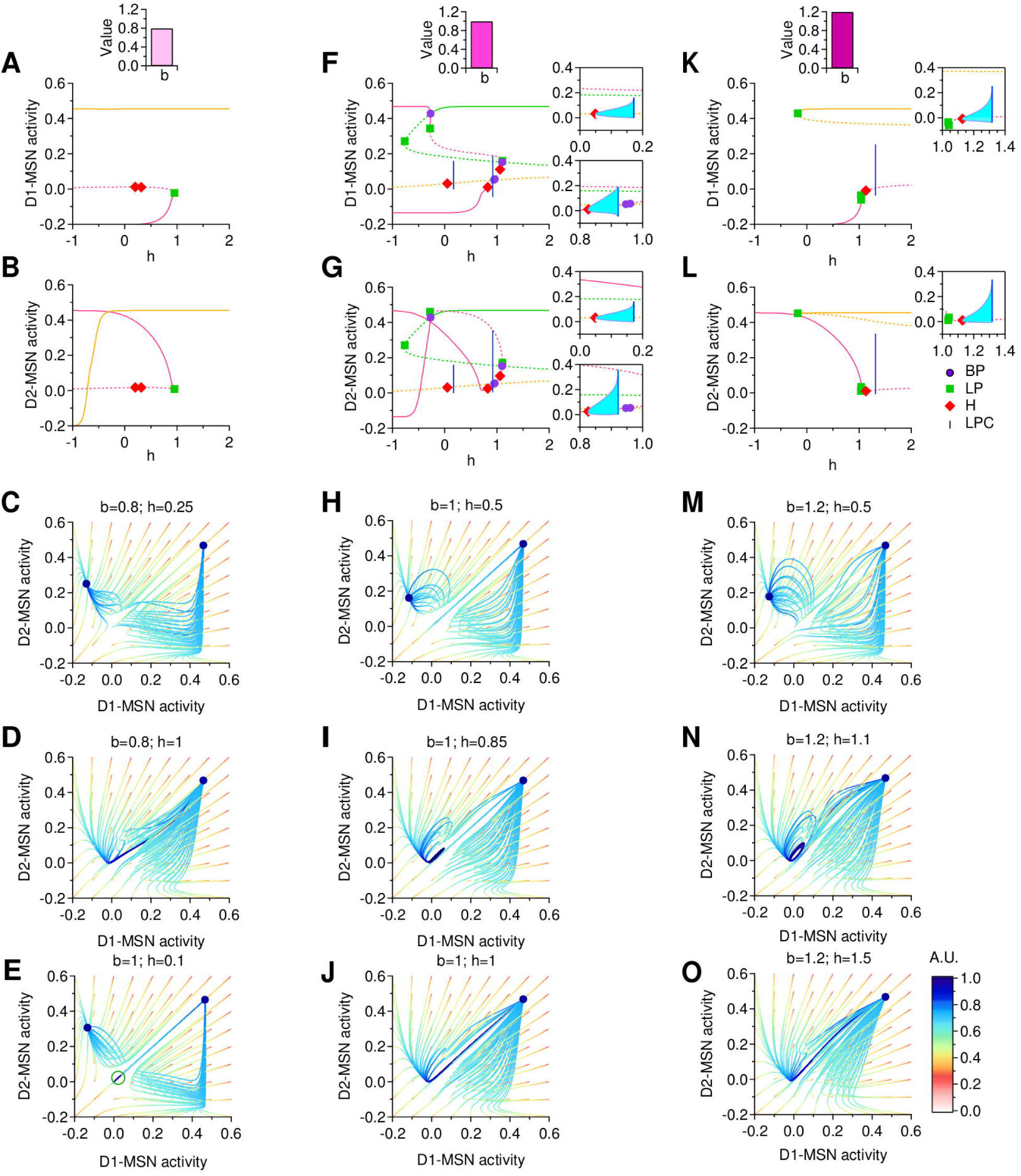
Dynamic transitions for *b* = 0.8, *b* = 1 and *b* = 1.2. *(A*-*B)*, *Bifurcation diagrams with respect to h*, *for D1*- *and D2*-*MSNs*, *when b* = 0.8*. (C*-*D)*, *Trajectories in a* (*D*1, *D*2) *phase space slice for b* = 0.8 *and h* = 0.25 *and h* = 1*. (E) As in (C*-*D)*, *for b* = 1 *and h* = 0.1*. (F*-*G) As in (A*-*B)*, *for b* = 1*. (H*-*J) As in (C*-*D)*, *for b* = 1 *and for different values of h. (K*-*L) As in (A*-*B) for b* = 1.2*. (M*-*O) As in (C*-*D)*, *for b* = 1.2 *and for different values of h. Fixed parameters: c*_*e*_0__ = *c*_*i*_0__ = 20.

### Effect of local and coupled changes in the E/I of D1- and D2 -MSNs

As a last step in our analysis, we ask how simultaneous, coupled changes of both excitation and inhibition onto D1- and D2-MSNs can alter the activity level of these cells, using the following system of equations:

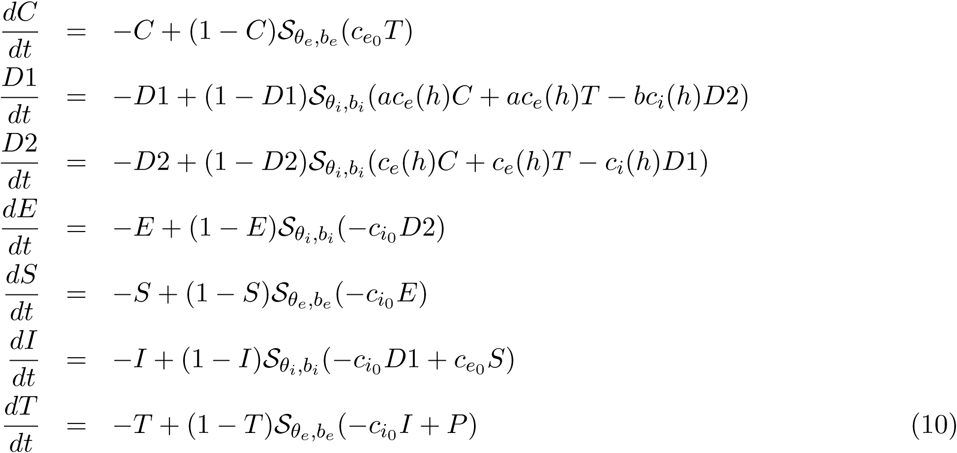

We use the parameter *h* to simultaneously change the E/I in both D1- and D2-MSNs (i.e. larger values of *h* correspond to an increasing excitation and decrease in inhibition onto D1- and D2-MSNs). The parameters *a* and *b* allow us to scale the relative excitation and inhibition level onto D1 and D2-MSNs. Fig 10 shows how the system can be pushed through a sequence of bifurcations and a succession of regimes by increasing the parameter *h*, for two instances of (*a*, *b*), shown side by side. In Fig 10A-F, *a* = 1 and *b* = 1. In Fig 10G-L, *a* = 1 and *b* = 1.2. Both sets of models show qualitatively similar results. Accordingly, for small values of *h*, the system resides in a bi-stable regime of differentially high/low activity for D1- and D2-MSNs. As *h* increases, the system shows stable out-of-phase oscillations until both D1- and D2-MSNs show high levels of activity (Fig 10A-F). Increasing *b* from *b* = 1 to *b* = 1.2 does not change the occurrence of these transitions, but causes subtler, yet important effects (Fig 10G-L). *First*, for the higher *b*, the transition between the bi-stability and mono-stability regimes for increasing values of *h* is interrupted by two LPs, which cause an abrupt transition in the equilibrium curves. During this transition, for 0.25 < *h* < 0.35, the system goes through a small window with a unique equilibrium, characterized by low activity in both D1 and D2-MSNs. This suggests that when inhibition decreases more in D1- than D2-MSNs, the system passes through a transient state of basal D1- and D2-MSN activity before reaching a mono-stability regime. *Second*, the stable oscillation window between the LPC and the H bifurcation points is prolonged. This is important, since the presence of oscillations prevents the system from converging to a state of steady hyperactivity for D1- and D2-MSNs.

**Fig 10.**
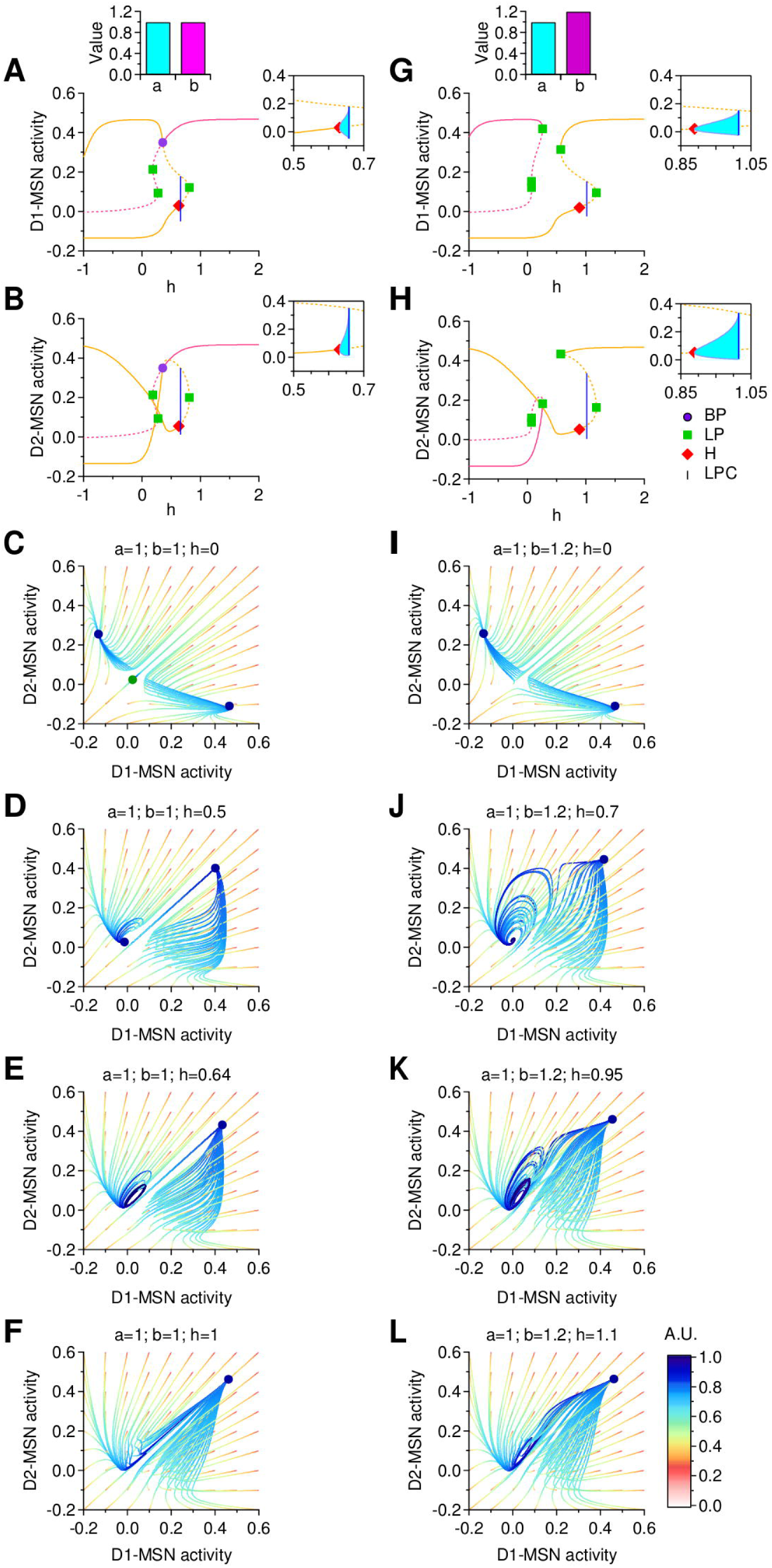
Dynamic transitions for (*a* = 1, *b* = 1) and (*a* = 1, *b* = 1.2). *(A*-*B) Bifurcations with respect to h*, *for D1*- *and D2*-*MSNs. (C*-*F) Trajectories in a* (*D*1, *D*2) *phase space slice for (top to bottom): h* = 0; *h* = 0.5; *h* = 0.64; *h* = 1*. (G*-*H) Bifurcations with respect to h*, *for D1*- *and D2*-*MSNs. (I*-*L) Trajectories in a* (*D*1, *D*2) *phase space slice for (top to bottom): h* = 0*; h* = 0.7*; h* = 0.95*; h* = 1.1*. Fixed parameters: c*_*e*_0__ = *c*_*i*_0__ = 20.

## Discussion

The goal of this study is to detemine how global and local changes in excitation and inhibition in distinct domains of the CSTC pathway affect the activity of this brain circuit and of specific classes of striatal neurons. The two major pieces of evidence that motivate our study are: *(1)* the CSTC pathway controls important physiological functions, like movement execution; *(2)* the CSTC pathway is a major site of synaptic dysfunction in neurospychiatric disorders like OCD and Tourette’s syndrome, which are characterized by the repeated execution of involuntary movements [22, 9]. In this pathway, cortical and thalamic neurons projecting to the striatum form glutamatergic synapses onto D1- and D2-MSNs. These cells form local inhibitory GABAergic synapses onto each other and send long-projection GABAergic afferents to the substantia nigra pars reticulata (SNr) and the internal capsule of the globus pallidus (GPi) via two anatomically distinct pathways (the direct and indirect pathways, respectively). Information from D1- and D2-MSNs is ultimately relayed to the thalamus and the motor cortex [23]. Coordinated activation of D1- and D2-MSNs is crucial for action selection and inhibition of unwanted behaviors as well as to control the overall level of activity in the CSTC pathway [24, 25]. Accordingly, reduced activity of D1-MSNs and increased activity of D2-MSNs generate a powerful inhibition of the motor cortex. In contrast, increased activity of D1-MSNs and reduced activity of D2-MSNs promote motor cortex activity and movement execution, which is thought to contribute to the motor symptoms of OCD [26, 27, 28, 29]. Consistent with these findings, clinical sudies in humans indicate that patients with OCD show functional hyperactivity in the CSTC pathway. These findings are largely based on the use of positron emission tomography (PET) and single photon emission CT (SPECT) [30, 31, 32, 33, 34, 35] and on resting-state functional magnetic resonance imaging (rs-fMRI) [36, 37, 38, 39], all of which lack single-cell resolution. A recent connectomics study based on the combined use of magnetic resonance scans and deterministic fiber tracking based on diffusion tensor imaging data indicate that these functional alterations in the CSTC pathway are associated with structural alterations in the *connectivity strength* between the orbito-frontal cortex and the striatum, two important nodes of the CSTC pathway [40]. What is not known is whether these functional and structural abnormalities in the CSTC pathway can be attributed to changes in excitatory and/or inhibitory transmission onto MSNs.

In this work, we model the activity of the CSTC pathway using a system of seven non-linear equations, to provide a mathematical description of the currently known wiring diagram of the CSTC pathway. Our modeling approach, originally proposed by Wilson and Cowan [12], allows us to describe the mean field activity in neural populations, not the firing rate of individual cells (consistently with the level of resolution of neuroimaging works). According to this definition, a given population firing rate can be due to stereotyped firing rate in all neurons or to the presence of a small number of high firing neurons. The spontaneous activity of medium spiny neurons in the striatum involves recurring shifts in membrane potential from a hyperpolarized (down) state to a depolarized (up) state accompanied by irregular spikes at low frequency and burst firing [43, 44, 45, 46]. This is different from the activity of other types of neurons (e.g. dopaminergic neurons in the ventral tegmental area of the midbrain [13], which exhibit ongoing (tonic) firing activity interleaved by brief periods of bursting (phasic) activity [42]. In this context, the Wilson-Cowan type model used here allows us to accurately capture the temporal evolution of the mean level of activity of distinct neuronal populations without adding further information on the specific firing activity of individual cells.

Although some works suggest that glutamatergic synaptic dysfunction contributes to the etiology of OCD [54], others indicate that changes in GABAergic inhibition disrupt the activity of the CSTC pathway and lead to the onset of OCD-like behaviors [11, 55, 56, 57]. Our network analysis sheds light on this conundrum. They indicate that changes in excitation or inhibitiion can both cause an increased activity in D1- and reduced activity in D2-MSNs (see red cells in Figure 11). In particular, this pattern of activity can be generated by: *(1)* increasing excitation on both D1- and D2-MSNs (Fig 4); *(2)* varying excitation onto D1-MSNs at low levels of excitation of D2-MSNs (Fig 5, 8); *(3)* decreasing inhibition of D1-MSNs when excitation of both MSNs is low (Fig 10). Varying GABAergic inhibition onto MSNs exerts a more powerful control of the oscillatory activity (than the overal level of activity) of D1 and D2-MSNs. Power spectra analysis from striatal local field potentials and electrocorticograms highlight the presence of two major types of spontaneous oscillatory activities in the low (2-9 Hz; *δ*, *θ*) and high frequency range (30-80 Hz; *γ*) [58]. A dynamic progression in the dominant mode of oscillatory activity is thought to occur in the striatum when learning (ventral striatum) and when performing a specific motor repertoire (dorsal striatum) [59, 60, 61]. According to these findings, the oscillatory activity in the ventral striatum evolves from transient bouts in the *γ* range to transient bouts in the *β* range (15-28 Hz) as a specific movement is consolidated into a habit through learning [61]. The ventral striatum is thought to direct the activity of the dorsal striatum through corresponding changes in its oscillatory activity [59]. For this reason, changes in the relative inhibition of D1- and D2-MSNs provide a valuable candidate mechanism that contributes to disrupted habit formation and movement execution in OCD.

In our analysis, we point out that there are wide parameter ranges that confer bi-stability to the system. In particular, the asymptotic trajectory of the system depends crucially on the positions of its initial state. In most cases, the hyperspace *D*1 = *D*2 delimits a boundary between two different attracting regimes: one with high D1- and low D2-MSN activity, a second with low D1- and high D2-MSN activity (Fig 5, 8-10). A small initial bias in the activity of either D1- or D2-MSNs would lead the system to one of these attractors. In many instances, the hyperspace *D*1 = *D*2 contains a saddle equilibrium or a saddle cycle. Only initial conditions in which the activity of D1-MSNs is identical to that of D2-MSNs would maintain the system in this hyperspace. However, regimes with perfectly identical D1- and D2-MSN activity are extremely unlikely to be reached, although other factors that have not included in this model (e.g. spike timing) could make this regime more likely to be maintained.

Taken together, these findings allow us to analyze the effects of small perturbations on the activity of D1- and D2-MSNs and of the entire CSTC pathway. They highlight that local inhibition, among all other factors, can powerfully regulate the activity of the entire system, possibly contributing to the hyperactivity patterns observed in patients with OCD.

Figure 11: **Summary table of all observed effects when changing different parameters.**

## Acknowledgments

This work was supported by: SUNY New Paltz Provost Challenge Grant (A.R.R.); National Science Foundaton (IOS 1655365), SUNY Albany and SUNY Albany Research Foundation (A.S.).

## Author contributions

ARR, AS conceived research; ARR, JH, CK performed research; ARR, AS wrote the manuscript.

## Additional information

The authors declare no competing financial interests.

**Table 1.**

| Parameters | Effect | Reference | Parameters | Effect | Reference |
| --- | --- | --- | --- | --- | --- |
| $\downarrow c_e, \downarrow c_i$ | $\downarrow D1, D2 (\downarrow C, T)$ | Fig 2 | $\Delta h, a=1$ | $\uparrow D1; \downarrow D2$ | Fig 8 |
| | oscillations | Fig 3 | | $\downarrow D1; \uparrow D2$ | Fig 8 |
| $\uparrow c_e, \uparrow c_i$ | $\uparrow D1, D2 (\uparrow C, T)$ | Fig 2 | | oscillations | Fig 8 |
| | oscillations | Fig 3 | $\Delta h, a=1.2$ | $\uparrow D1; \downarrow D2$ | Fig 8 |
| $\downarrow c_e, \uparrow c_i$ | $\downarrow D1, D2 (\downarrow C; \uparrow T)$ | Fig 2 | | $\downarrow D1; \uparrow D2$ | Fig 8 |
| | oscillations | Fig 3 | $\Delta h, b=0.8$ | $\uparrow D1, D2$ | Fig 9 |
| $\uparrow c_e, \downarrow c_i$ | $\uparrow D1, D2 (\uparrow C, T)$ | Fig 2 | | $\downarrow D1; \uparrow D2$ | Fig 9 |
| | oscillations | Fig 3 | $\Delta h, b=1$ | $\uparrow D1, D2$ | Fig 9 |
| $\Delta c_e'$ | $\uparrow D1; \downarrow D2 (\uparrow C, T)$ | Fig 4 | | $\downarrow D1, D2$ | Fig 9 |
| | $\downarrow D1; \uparrow D2 (\downarrow C, T)$ | Fig 4 | | oscillations | Fig 9 |
| $\Delta c_i'$ | $\uparrow D1, D2 (\uparrow C, T)$ | Fig 4 | $\Delta h, b=1.2$ | $\uparrow D1, D2$ | Fig 9 |
| | $\downarrow D1; \uparrow D2 (\downarrow C, T)$ | Fig 4 | | $\downarrow D1; \uparrow D2$ | Fig 9 |
| $\Delta c_{e1}, \downarrow c_{e2}$ | $\uparrow D1; \downarrow D2$ | Fig 5 | | oscillations | Fig 9 |
| | $\downarrow D1; \uparrow D2$ | Fig 5 | $\Delta h, a=1, b=1$ | $\uparrow D1, D2$ | Fig 10 |
| $\Delta c_{e1}, \uparrow c_{e2}$ | $\uparrow D1, D2$ | Fig 5 | | $\downarrow D1, D2$ | Fig 10 |
| | $\downarrow D1; \uparrow D2$ | Fig 5 | | oscillations | Fig 10 |
| | oscillations | Fig 5 | $\Delta h, a=1, b=1.2$ | $\uparrow D1; \downarrow D2$ | Fig 10 |
| $\Delta c_{i1}, \downarrow c_{i2}$ | $\uparrow D1, D2$ | Fig 6 | | $\downarrow D1; \uparrow D2$ | Fig 10 |
| | $\downarrow D1, D2$ | Fig 6 | | oscillations | Fig 10 |
|  | oscillations | Fig 6 |  |  |  |
| $\Delta c_{i1}, \uparrow c_{i2}$ | $\uparrow D1, D2$ | Fig 6 | | | |
| | $\downarrow D1; \uparrow D2$ | Fig 6 | | | |
|  | oscillations | Fig 6 |  |  |  |

## References

[1] Ann M Graybiel, Toshihiko Aosaki, Alice W Flaherty, Minoru Kimura, et al. The basal ganglia and adaptive motor control. SCIENCE-NEW YORK THEN WASHINGTON-, pages 1826–1826, 1994.

[2] Joshua D Berke, Murat Okatan, Jennifer Skurski, and Howard B Eichenbaum. Oscillatory entrainment of striatal neurons in freely moving rats. Neuron, 43(6):883–896, 2004.

[3] Norman M White. Mnemonic functions of the basal ganglia. Current opinion in neurobiology, 7(2):164–169, 1997.

[4] Edward A Stern, Anthony E Kincaid, and Charles J Wilson. Spontaneous subthreshold membrane potential fluctuations and action potential variability of rat corticostriatal and striatal neurons in vivo. Journal of Neurophysiology, 77(4):1697–1715, 1997.

[5] Charles R Gerfen and Kristen A Keefe. Neostriatal dopamine receptors. Trends in Neurosciences, 17(1):2–3, 1994.

[6] Guohong Cui, et al. Concurrent activation of striatal direct and indirect pathways during action initiation. Nature, 494(7436):238–242, 2013.

[7] Alexxai V Kravitz, Lynne D Tye, and Anatol C Kreitzer. Distinct roles for direct and indirect pathway striatal neurons in reinforcement. Nature neuroscience, 15(6):816–818, 2012.

[8] Sanjaya Saxena and Scott L Rauch. Functional neuroimaging and the neuroanatomy of obsessive-compulsive disorder. Psychiatric Clinics of North America, 23(3):563–586, 2000.

[9] Jonathan T Ting and Guoping Feng. Neurobiology of obsessive–compulsive disorder: insights into neural circuitry dysfunction through mouse genetics. Current opinion in neurobiology, 21(6):842–848., 2011.

[10] Susanne E Ahmari, et al. Repeated cortico-striatal stimulation generates persistent ocd-like behavior. Science, 340(6137):1234–1239, 2013.

[11] Eric Burguiére, Patrícia Monteiro, Guoping Feng, and Ann M Graybiel. Optogenetic stimulation of lateral orbitofronto-striatal pathway suppresses compulsive behaviors. Science, 340(6137):1243–1246, 2013.

[12] Hugh R Wilson and Jack D Cowan. Excitatory and inhibitory interactions in localized populations of model neurons. Biophysical journal, 12(1):1, 1972.

[13] Anca R Radulescu. Mechanisms explaining transitions between tonic and phasic firing in neuronal populations as predicted by a low dimensional firing rate model. PloS one. 5(9): e12695; 2010.

[14] Galina N Borisyuk, Roman M Borisyuk, Alexander I Khibnik, and Dirk Roose. Dynamics and bifurcations of two coupled neural oscillators with different connection types. Bulletin of mathematical biology, 57(6):809–840, 1995.

[15] Alain Destexhe, and Terrence Sejnowski. The Wilson–Cowan model, 36 years later. Biological cybernetics, 101(1): 1–2, 2009.

[16] Jack Cowan, Jeremy Neuman, and Wim van Drongelen. Wilson–Cowan equations for neocortical dynamics. The Journal of Mathematical Neuroscience. 6(1): 1, 2016.

[17] Vladimir Shusterman, and William C Troy. From baseline to epileptiform activity: a path to synchronized rhythmicity in large-scale neural networks. Physical Review E.

[18] Zachary P Kilpatrick, and Paul C Bressloff. Effects of synaptic depression and adaptation on spatiotemporal dynamics of an excitatory neuronal network. Physica D: Nonlinear Phenomena. 239(9): 547–560; 2010.

[19] Anca Răadulescu and Sergio Verduzco-Flores. Nonlinear network dynamics under perturbations of the underlying graph. Chaos: An Interdisciplinary Journal of Nonlinear Science, 25(1), 2015.

[20] Annick Dhooge, Willy Govaerts, and Yu A Kuznetsov. Matcont: a matlab package for numerical bifurcation analysis of odes. ACM SIGSAM Bulletin, 38(1):21–22, 2004.

[21] Annick Dhooge, Willy Govaerts, Yu A Kuznetsov, Hge Meijer, and Bart Sautois. New features of the software matcont for bifurcation analysis of dynamical systems. Mathematical and Computer Modelling of Dynamical Systems, 14(2):147–175, 2008.

[22] Jonathan T Ting and Guoping Feng. Glutamatergic synaptic dysfunction and obsessive-compulsive disorder. Current chemical genomics, 2(1), 2008.

[23] Ann M Graybiel. The basal ganglia. Current Biology, 10(14):R509–R511, 2000.

[24] JB Penney and AB Young. Speculations on the functional anatomy of basal ganglia disorders. Annual review of neuroscience, 6(1):73–94, 1983.

[25] Adam Ponzi and Jeffery R Wickens. Optimal balance of the striatal medium spiny neuron network. PLoS Comput Biol, 9(4):e1002954, 2013.

[26] Roger L Albin, Anne B Young, and John B Penney. The functional anatomy of basal ganglia disorders. Trends in neurosciences, 12(10):366–375, 1989.

[27] Garrett E Alexander and Michael D Crutcher. Functional architecture of basal ganglia circuits: neural substrates of parallel processing. Trends in neurosciences, 13(7):266–271, 1990.

[28] Peter Jenner. The rationale for the use of dopamine agonists in parkinson’s disease. Neurology, 45(3 Suppl 3):S6–S12, 1995.

[29] K Takakusaki, Kazuya Saitoh, H Harada, and M TakakusakiKashiwayanagi. Role of basal ganglia–brainstem pathways in the control of motor behaviors. Neuroscience research, 50(2):137–151, 2004.

[30] Lewis R Baxter, et al. Local cerebral glucose metabolic rates in obsessive-compulsive disorder: a comparison with rates in unipolar depression and in normal controls. Archives of General Psychiatry, 44(3):211–218, 1987.

[31] John C Mazziotta, Michael E Phelps, and Jorg J Pahl. Cerebral glucose metabolic rates in nondepressed patients with obsessive compulsive disorder. Am J Psychiatry, 145:1560–3, 1988.

[32] Susan E Swedo, et al. Cerebral glucose metabolism in childhood-onset obsessive-compulsive disorder. Archives of General Psychiatry, 46(6):518–523, 1989.

[33] GV Sawle, NF Hymas, AJ Lees, and RSJ Frackowiak. Obsessional slowness. Brain, 114(5):2191–2202, 1991.

[34] Scott L Rauch, et al. Regional cerebral blood flow measured during symptom provocation in obsessive-compulsive disorder using oxygen 15labeled carbon dioxide and positron emission tomography. Archives of General Psychiatry, 51(1):62–70, 1994.

[35] PK McGuire, et al. Functional anatomy of obsessive-compulsive phenomena. The British Journal of Psychiatry, 164(4):459–468, 1994.

[36] Tijiang Zhang, Yanchun Yang, Bin Li, Qiang Yue, and Yufeng Zang. Abnormal small-world architecture of top-down control networks in obsessive-compulsive disorder. Journal of psychiatry & neuroscience: JPN, 36(1):23, 2011.

[37] Da-Jung Shin, et al. The effects of pharmacological treatment on functional brain connectome in obsessive-compulsive disorder. Biological psychiatry, 75(8):606–614, 2014.

[38] Jing-Ming Hou, et al. Resting-state functional connectivity abnormalities in patients with obsessive-compulsive disorder and their healthy first-degree relatives. Journal of psychiatry & neuroscience: JPN, 39(5):304, 2014.

[39] Martin Göttlich, Ulrike M Krämer, Andreas Kordon, Fritz Hohagen, and Bartosz Zurowski. Decreased limbic and increased fronto-parietal connectivity in unmedicated patients with obsessive-compulsive disorder. Human brain mapping, 35(11):5617–5632, 2014.

[40] TJ Reess, et al. Connectomics-based structural network alterations in obsessive-compulsive disorder. Translational Psychiatry, 6(9):e882, 2016.

[41] Edward A Stern, Anthony E Kincaid, and Charles J Wilson. Spontaneous subthreshold membrane potential fluctuations and action potential variability of rat corticostriatal and striatal neurons in vivo. Journal of Neurophysiology, 77(4):1697–1715, 1997.

[42] Sabine Krabbe, et al. Increased dopamine d2 receptor activity in the striatum alters the firing pattern of dopamine neurons in the ventral tegmental area. Proceedings of the National Academy of Sciences, 112(12):E1498–E1506, 2015.

[43] Dietmar Plenz and Stephen T Kitai. Up and down states in striatal medium spiny neurons simultaneously recorded with spontaneous activity in fast-spiking interneurons studied in cortex–striatum–substantia nigra organotypic cultures. Journal of Neuroscience, 18(1):266–283, 1998.

[44] J Wayne Aldridge and Sid Gilman. The temporal structure of spike trains in the primate basal ganglia: afferent regulation of bursting demonstrated with precentral cerebral cortical ablation. Brain research, 543(1):123–138, 1991.

[45] Charles J Wilson and Philip M Groves. Spontaneous firing patterns of identified spiny neurons in the rat neostriatum. Brain research, 220(1):67–80, 1981.

[46] Charles J Wilson. The generation of natural firing patterns in neostriatal neurons. Progress in brain research, 99:277–297, 1993.

[47] Edward A Stern, Dieter Jaeger, and Charles J Wilson. Membrane potential synchrony of simultaneously recorded striatal spiny neurons in vivo. Nature, 394(6692):475–478, 1998.

[48] D Plenz and A Aertsen. Neural dynamics in cortex-striatum co-culturesii. spatiotemporal characteristics of neuronal activity. Neuroscience, 70(4):893–924, 1996.

[49] Charles J Wilson and Yasuo Kawaguchi. The origins of two-state spontaneous membrane potential fluctuations of neostriatal spiny neurons. Journal of neuroscience, 16(7):2397–2410, 1996.

[50] H Kita. Glutamatergic and gabaergic postsynaptic responses of striatal spiny neurons to intrastriatal and cortical stimulation recorded in slice preparations. Neuroscience, 70(4):925–940, 1996.

[51] BD Bennett and JP Bolam. Synaptic input and output of parvalbumin-immunoreactive neurons in the neostriatum of the rat. Neuroscience, 62(3):707–719, 1994.

[52] H Kita, T Kosaka, and CW Heizmann. Parvalbumin-immunoreactive neurons in the rat neostriatum: a light and electron microscopic study. Brain research, 536(1):1–15, 1990.

[53] HB Parthasarathy and AM Graybiel. Cortically driven immediate-early gene expression reflects modular influence of sensorimotor cortex on identified striatal neurons in the squirrel monkey. Journal of Neuroscience, 17(7):2477–2491, 1997.

[54] Eric J Nordstrom, Katie C Bittner, Michael J McGrath, Clinton R Parks, and Frank H Burton. “hyperglutamatergic cortico-striato-thalamo-cortical circuit” breaker drugs alleviate tics in a transgenic circuit model of tourette’s syndrome. Brain research, 1629:38–53, 2015.

[55] Paul SA Kalanithi, et al. Altered parvalbumin-positive neuron distribution in basal ganglia of individuals with tourette syndrome. Proceedings of the National Academy of Sciences of the United States of America, 102(37):13307–13312, 2005.

[56] Yuko Kataoka, et al. Decreased number of parvalbumin and cholinergic interneurons in the striatum of individuals with tourette syndrome. Journal of Comparative Neurology, 518(3):277–291, 2010.

[57] Meiyu Xu, et al. Targeted ablation of cholinergic interneurons in the dorsolateral striatum produces behavioral manifestations of tourette syndrome. Proceedings of the National Academy of Sciences, 112(3):893–898, 2015.

[58] Andrew Sharott, et al. Different subtypes of striatal neurons are selectively modulated by cortical oscillations. The Journal of Neuroscience, 29(14):4571–4585, 2009.

[59] Hisham E Atallah, Dan Lopez-Paniagua, Jerry W Rudy, and Randall C O’Reilly. Separate neural substrates for skill learning and performance in the ventral and dorsal striatum. Nature neuroscience, 10(1):126–131, 2007.

[60] Pepe J Hernandez, Kenneth Sadeghian, and Ann E Kelley. Early consolidation of instrumental learning requires protein synthesis in the nucleus accumbens. Nature neuroscience, 5(12):1327–1331, 2002.

[61] Mark W Howe, Hisham E Atallah, Andrew McCool, Daniel J Gibson, and Ann M Graybiel. Habit learning is associated with major shifts in frequencies of oscillatory activity and synchronized spike firing in striatum. Proceedings of the National Academy of Sciences, 108(40):16801–16806, 2011.

